# Enzyme-mediated alkynylation enables transcriptome-wide identification of pseudouridine modifications

**DOI:** 10.1101/2023.06.18.545436

**Authors:** Yuru Wang, Lisheng Zhang, Wen Zhang, Boyang Gao, Chang Ye, Qing Dai, Ke Wang, Minkui Luo, Tao Pan, Chuan He

## Abstract

Pseudouridine (Ψ) is one of the most abundant chemical modifications that exists in various types of RNA species and is known to play important roles in RNA function. The advances in studies of Ψ in less abundant messenger RNA species have been hindered by a lack of suitable methods to precisely and sensitively map their distributions. Here we show that a methyltransferase from *Methanocaldococcus jannaschii* can label RNA Ψ efficiently and specifically with various functional groups, both in isolated RNA and inside cells. We leveraged this enzymatic labeling strategy to develop ELAP-seq as a facile method to enrich Ψ-modified transcripts for the detection of Ψ modifications at single base resolution with high sensitivity and low background. Using this method, we identified over 10, 000 candidate Ψ sites from human transcripts, which provides new insights into Ψ biosynthesis and function. Our study provides a chemical biology method that specifically labels Ψ for its detection and functional alteration.

## Introduction

Chemical modifications of RNA are critical to most RNA functions^1^. Among the over 150 types of RNA chemical modifications, pseuoduridine (Ψ) is one of the most abundant and exists in multiple types of RNA species, including ribosome RNA (rRNA), transfer RNAs (tRNA), small nucleolar RNAs (snoRNA) and messenger RNAs (mRNAs)^2, 3^. Past studies have shown that the presence of Ψ in RNA can alter RNA properties including structure, stability, alternative splicing and translation^4–7^. As a consequence, Ψ modifications can have a profound effect on cellular activities such as cell proliferation and stem cell differentiation^8, 9^. Ψ has also been found to widely exist in virus RNAs and may play roles in viral infection and anti-viral defense pathways^10^.

Ψ is formed via the isomerization of uridine, a reaction catalyzed either by various stand-alone Ψ synthases (PUS) or the DKC1 complex guided by box H/ACA small nucleolar RNAs (snoRNAs)^3, 11^. The enzymatic cores of the PUS enzymes are conserved, and their malfunctions are often associated with human diseases, including autoimmune digestive disorders and intellectual disability^12–14^. However, how the loss of Ψ contributes to human diseases has yet to be fully elucidated.

Ψ and its derivative N^1^-methylpseudouridine (m^1^Ψ) are among the most critical nucleoside substitutions that notably promote efficacies of mRNA vaccines^15^. These modifications prevent mRNA vaccines from triggering unwanted innate immune response and promote translation of the modified mRNA. The therapeutic potentials of Ψ and m^1^Ψ have inspired enormous interests to investigate their natural presence and biological functions in cells. A more comprehensive understanding of the pathways by which these modified nucleosides affect mRNA stability, translation and immunity may facilitate additional therapeutic developments.

While Ψ has been studied in the context of abundant rRNA, tRNAs and snoRNAs for decades, the significance of Ψ in less abundant mRNAs is less well understood. This is limited by a lack of suitable methods to detect Ψ modifications sensitively and precisely in mRNA species. Pseudouridine has the same base pairing property as uridine, making it impossible to directly distinguish this modification from uridine based on Sanger or next-generation sequencing. Past methods to detect Ψ modification sites rely on chemical treatments to convert Ψ to forms generating stop or deletion signatures at or near the modification sites during reverse transcription^16–20^. For examples, recently developed methods BID-seq and PRAISE rely on bisulfite treatment and allow quantitative mapping of Ψ transcriptome-wide^16, 20–22^. While accurate and selective, these methods do not involve enrichment, therefore require deep sequencing in order to detect and quantify deletion signature induced by a Ψ-bisulfite adduct. Lowly expressed genes and less modified sites could be missed when sequencing depth is not deep enough. Enrichment based method has been developed based on the CMC addition reaction; however, non-specific addition reaction to other nucleobases could lead to a non-negligible rate of false positives which can be further exaggerated under the condition of enrichment^19^. Direct Ψ detection methods based on nanopore sequencing technology have also been developed; however, these methods are currently limited by their relatively low resolution, requirement for large amount of RNAs, and complicated data analysis pipelines^23, 24^.

LC-MS/MS reported Ψ abundance in mRNA close to *N*^6^-methyladenosine (m^6^A, 0.2-0.4% in HeLa and HEK 293T cell lines)^19^; however, currently reported numbers of Ψ modification sites in routinely used lab cell lines are still far from its actual abundance, suggesting more sites are to be discovered. Thus, new methods using independent strategies i is highly desired to comprehensively identify modification sites as well as to cross-validate previously reported ones. Importantly, currently existing labeling strategies cause RNA degradation and are only limited to isolated RNAs. There is no feasible way to efficiently and specifically label Ψ in situ and inside cells so far.

To provide additional approaches for both *in vitro* and potential cell-based manipulation of Ψ, we turned to nature for inspirations on new labeling strategies for Ψ. Nature has evolved a plethora of proteins and enzymes that selectively recognize and act on RNA modifications; exploiting these biomolecules offers a promising new direction for modulation of different RNA modifications^25^. After examining several enzymes for *in vitro* reactivity, we focused on a methyltransferase from *Methanocaldococcus jannaschii*^26^. We found that this enzyme can install a methyl or propargyl group specifically onto the N1 position of Ψ *in vitro* in various sequence contexts. This discovery offers convenient and mild one-step reaction to specifically label Ψ under physiological condition, and can be used to modulate Ψ inside human cells, making it highly promising and versatile strategy to detect and modulate Ψ.

To showcase the utilize of this selective labeling strategy, we firstly developed a method that specifically enriches Ψ-containing transcripts for transcriptome-wide Ψ profiling. We validated the method by applying it to detect known Ψ modifications in rRNA, as well as successfully identify over 10,000 candidate Ψ modification sites in the transcriptomes of HeLa and HEK 293T cell lines, providing additional insights into features of Ψ distribution and function in human cells. Altogether, our study provides new tools that allow studies of Ψ modification both *in vitro* and in cells.

## Results

### A Ψ-specific methyltransferase that specifically labels Ψ modifications

Currently, there are limited labeling tools to investigate RNA Ψ modification *in vitro* and no such tools work inside cells. We decided to explore natural enzymes that specifically act on Ψ. Natural enzymes often provide useful starting points to develop biotechnologies because of their exceptional substrate specificity and catalytic efficiency^25^. One major difference in physiochemical features between uridine and Ψ lies in an extra hydrogen donor at the N1 position of the Ψ nucleobase (Figure 1a). We reasoned natural enzymes that specifically react at this position would facilitate selective labeling of Ψ over uridine.

**Figure 1.**
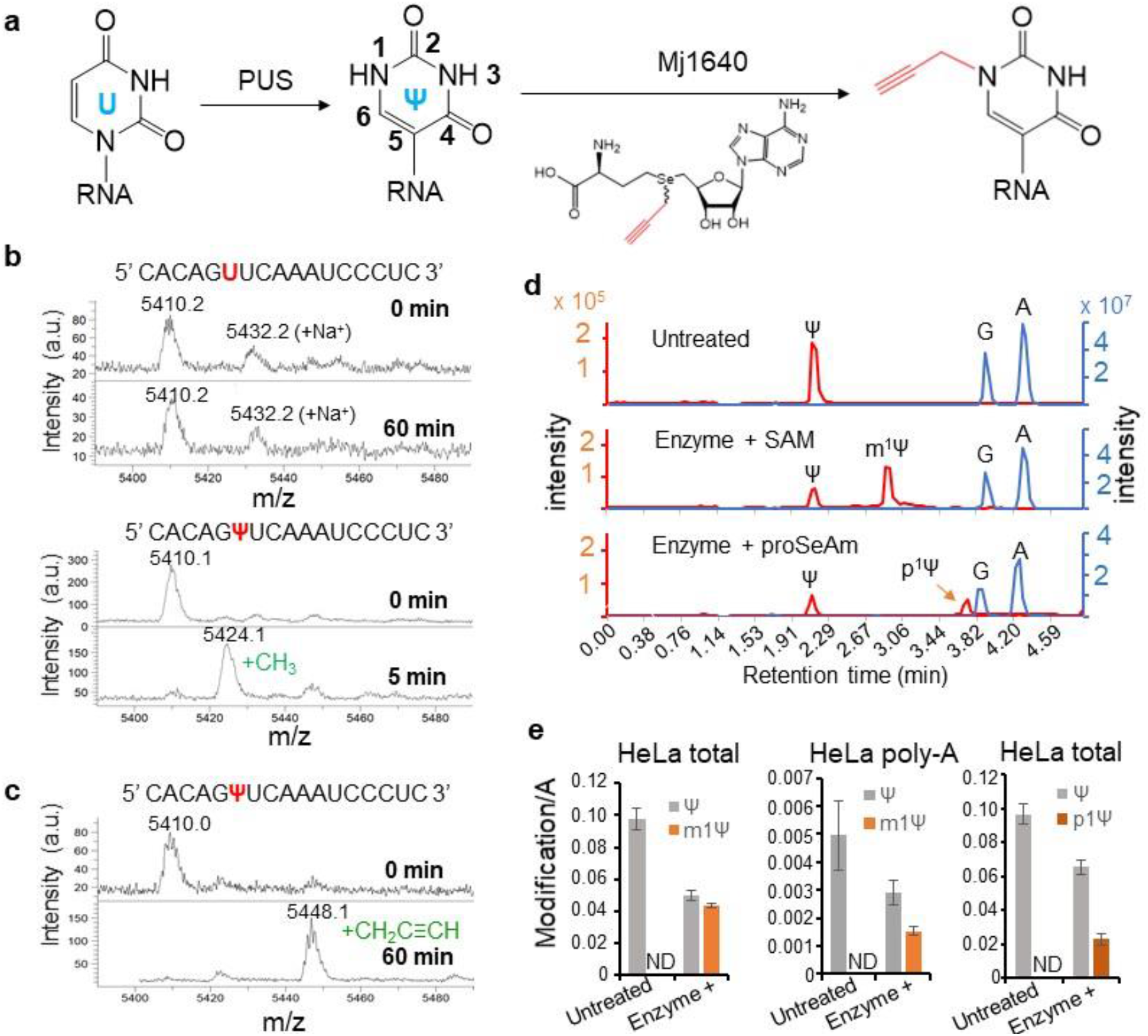
Methyltransferase Mj1640 can label Ψ specifically and promiscuously *in vitro*. **a**) Enzymatic conversions of uridine to Ψ and Ψ to *N*^1^-propargylΨ (p^1^Ψ). **b**) MALDI-TOF MS shows that Mj1640 specifically methylates Ψ but not U in synthetic RNA oligonucleotides. **c**) MALDI-TOF MS shows that Mj1640 can transfer a propargyl group onto Ψ in the synthetic RNA oligonucleotide by making use of Pro-SeAm co-factor. **d**) LC-MS/MS signals of nucleosides Ψ, *N*^1^-methylΨ (m^1^Ψ) and p^1^Ψ in digested total RNA with and without enzymatic treatment. **e**) Quantification of efficiencies of the enzymatic addition of the methyl or the propargyl group to Ψ in cellular total or poly-A RNA.

*N*^1^-methylpseudouridine (m^1^Ψ) is a modification naturally found in tRNA of most archaea and is formed via methylation at the N1 position of Ψ, a reaction catalyzed by SPOUT-class methyltransferases^27^. This modification plays important roles in tRNA structure and is also the most critical nucleobase substitution that vastly promotes the efficacy of the COVID-19 mRNA vaccine^28, 29^. We surveyed the SPOUT-class methyltransferases that catalyze this conversion and tested a few enzyme candidates. We focused our attention on the enzyme Mj1640 from *Methanocaldococcus jannaschii*, which is responsible for methylation of Ψ at position 54 in the T-arm of tRNAs (Figure S1a)^26^. Although the methylation was reported to occur naturally only on Ψ54 in tRNA, Wurm et al. reported that the enzymatic catalysis is highly efficient *in vitro* on a 17-mer RNA probe resembling the T-arm of *Methanocaldococcus* jannaschii tRNA^Trp^ at 65 °C with 100 mM K^+^ ion^26^. We reasoned that the relatively high reaction temperature may have disrupted the secondary structure of the 17-mer RNA probe, thus the enzyme might be more promiscuous for labeling Ψ independent of local RNA structure.

We proceed to express and purify the recombinant Mj1640 protein (Figure S1b-c), and tested its activity *in vitro* on the same 17-mer synthetic RNA oligonucleotide as in Wurm *et al.*^26^ and a control RNA with a U replacing the Ψ (Figure 1b and Figure S1e). The Ψ-RNA was methylated to > 90% in five minutes as evaluated by MALDI-TOF MS, whereas no reaction was observed for the U-RNA after over one hour incubation (Figure 1b), validating the excellent specificity of the reaction towards Ψ. Importantly, a synthetic 8-mer oligo containing a Ψ site was efficiently methylated (Figure S1d) whereas the control oligo with U was not, demonstrating that the enzymatic catalysis does not rely on RNA secondary structure. Sequence preference of this enzyme was also studied and will be shown later. We tested other Ψ specific methyltransferases including YbeA and MjNep1^30, 31^ but they either showed limited reactivity or had no reactivity *in vitro* (data not shown).

### Mj1640 can transfer allyl and propargyl groups to Ψ in RNA

Next, we tested whether the Mj1640 enzyme could use SAM derivatives to label Ψ sites with chemical groups that could be further modified. To our delight, Mj1640 efficiently utilized both allylic-SAM and propargyl-SeAm cofactors to label the 17-mer Ψ-RNA (> 80% in 1 hr) (Figure 1c and Figure S1e) but not the control U-RNA.

To evaluate the ability of the enzymatic reaction to label Ψ modifications in RNAs isolated from human cells, we assessed the labeling efficacy using quantitative LC-MS/MS. We detected a dramatic decrease of Ψ signal and an appearance of *N*^1^-methylpseudourisine (m^1^Ψ) or *N*^1^-porpargyl-pseudouridine (p^1^Ψ) signal after the enzymatic treatment using either SAM or propargyl-SeAm as the co-factor (Figure 1d and Figure S2a). With optimized protein purification workflow and enzymatic reaction condition (see the Methods section), the Ψ *in vitro* methylation ratio was around 50% for total RNA, and 33-48% and 50-60% for poly-A RNA extracted from HeLa and HEK293T cells, respectively, based on standard curves generated for each nucleoside (Figure 1e, Figure S2b-c). The 33-60% methylation ratio suggests that a considerable proportion of Ψ modification sites were successfully labeled. We confirmed the enzymatic addition of the propargyl group onto Ψ in cellular RNA via dot blot by adding on a biotin molecule via copper-catalyzed azide–alkyne cycloaddition (CuAAC or click) reaction (Figure S2g). We also quantified via LC-MS/MS that total RNA and poly-A RNA were labeled with the propargyl group at ∼25-30% of all Ψ (Figure 1e and Figure S2d-f). This slightly lower labeling percentage compared to methylation is likely due to a weaker binding affinity of the enzyme to propargyl-SeAm than to SAM, rather than a biased labeling. These results demonstrated the feasibility to label Ψ using Mj1640.

It is important to note that the enzymatic reaction condition is mild (pH 7.5 at 65 °C with the potassium ion), under which mRNAs remain largely intact (Figure S3a), highlighting the advantage of the enzymatic labeling strategy over chemical labeling strategies which typically cause RNA degradation. In addition, when expressed in human HEK 293T cells, the Mj1640 enzyme is functional to label Ψ in endogenous transcripts by converting Ψ to m^1^Ψ (Figure S3b-d). Altogether, our study presents a novel enzymatic strategy that can label Ψ both in isolated RNAs and inside cells.

### Development of the <u>E</u>nzymatic <u>La</u>beling and <u>P</u>ull-down for <u>Seq</u>uencing method (ELAP-seq) to detect Ψ on the transcriptome

To show an example of applying the enzymatic labeling strategy to studying Ψ, we developed a method coupling <u>E</u>nzymatic <u>La</u>beling and <u>P</u>ull-down for <u>Seq</u>uencing (ELAP-seq) to probe Ψ modification sites on the transcriptome (Figure 2a). The exclusive specificity of the enzymatic labeling towards Ψ is critical to ensure the enrichment of Ψ-modified RNA but not unmodified RNA. In this protocol, biological RNA such as total RNA or mRNA is first fragmented to 60-200 nucleotide, followed by the enzymatic treatment to label the Ψ sites with a propargyl group (Figure 2a). An aliquot of fragmented RNA prior to enzymatic treatment was used to directly prepare for sequencing library (input library), in order to control for signals from factors other than Ψ. The enzymatically labeled RNA is split into two parts, with one part undergone click reaction to add on a biotin molecule while the other part undergone a mock click reaction. Pull-down is then performed with magnetic beads coated with streptavidin to enrich RNA fragments containing Ψ modifications, followed by libraries construction and Illumina sequencing (Pull-down library).

**Figure 2.**
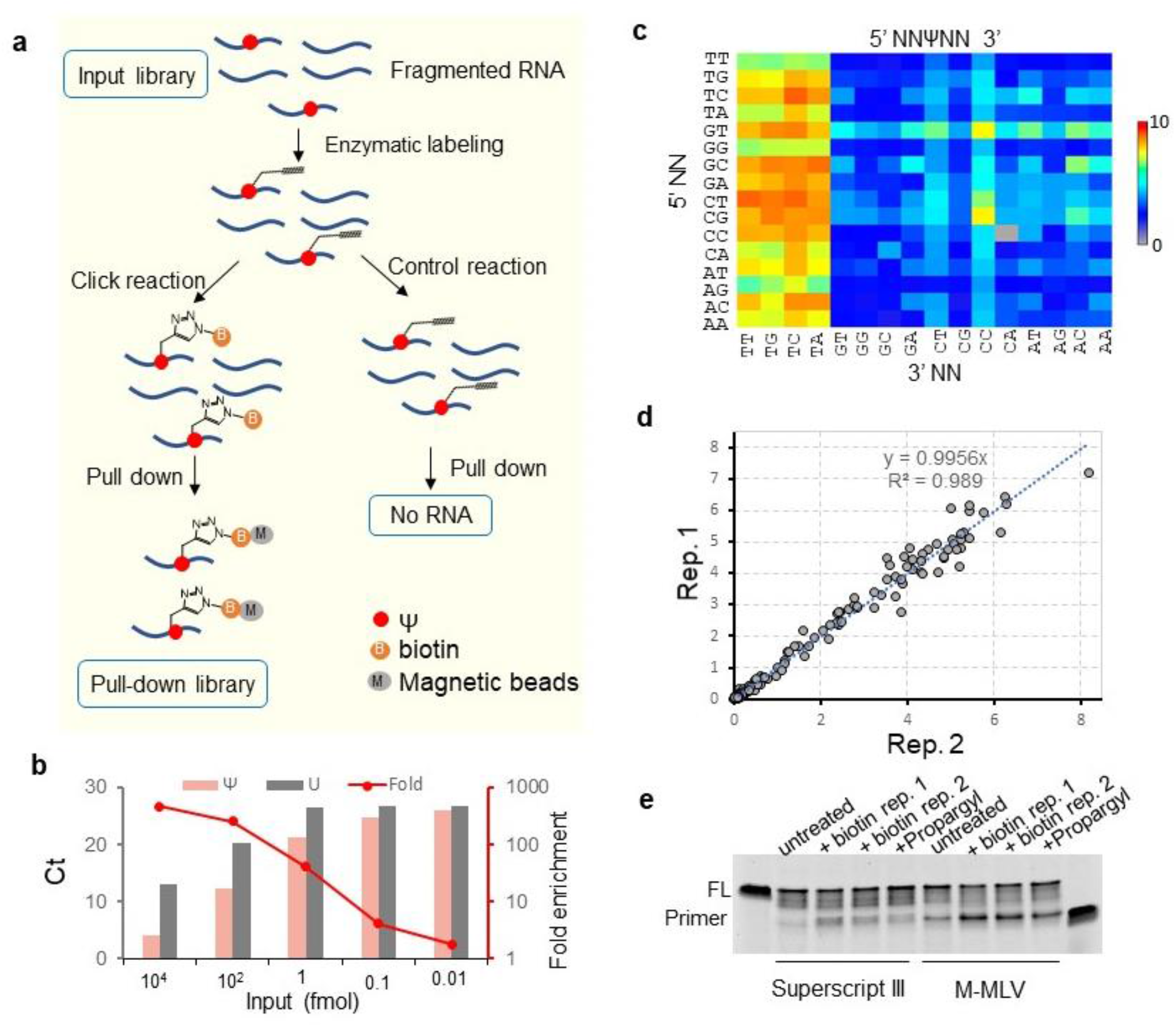
The development of the ELAP-seq method. **a**) ELAP-seq library preparation workflow. **b**) Pull-down and qPCR of RNA oligonucleotide with Ψ or U after enzymatic treatment and biotin addition reveal high signal-to-noise ratios. **c**) Heatmap shows various degrees of enrichment for synthetic oligonucleotides with 256 types of sequence motifs surrounding the Ψ modification (5’-NNΨNN) as evaluated by ELAP-seq. **d**) Correlation between two independent replicates on degrees of enrichment for synthetic RNA oligonucleotides with the 5’-NNΨNN motifs. Each dot represents one sequence motif. **e**) Superscript III and M-MLV RT under low dNTP concentration are arrested by *N*^1^-propargylΨ modification with (“+ biotin”) or without (“+ Propargyl”) clicking on a biotin molecule.

To evaluate the signal to noise ratio of the enzymatic labeling and pull-down strategy, we treated synthetic RNA oligonucleotides bearing either a Ψ or U in the center with enzymatic labeling and biotin addition sequentially. The RNA was then pulled down and quantified via qPCR. With optimization of the pull-down condition, the Ψ-RNA was found to be enriched for 4-450-folds relative to the U-RNA with input ranging from 0.01 fmol to 10 pmol (Figure 2b), demonstrating an excellent signal-to-noise ratio of the enzymatic labeling and pull-down strategy.

### Mj1640 preferentially labels Ψ in the 5’-ΨU-3’ sequence context

We then used ELAP-seq method to comprehensively evaluate the sequence context preference of the Mj1640 enzyme by building sequencing libraries from a mixture of 33-mer synthetic standard RNA oligonucleotides containing NNΨNN sequences (N means any nucleoside) which cover 256 sequence contexts surrounding Ψ. As expected, the control library yielded no sufficient material to produce a sequencing library in the end, highlighting a low background by the protocol. Interestingly, sequencing data from the Ψ-RNA showed that RNAs with 5’-ΨU-3’ context were preferentially enriched while marginal sequence preferences were observed at other nearby sites (Figure 2c). To further validate the observed sequence preference, we synthesized 10-mer RNA oligonucleotides containing degenerate A, U, C and G (denoted as N) at one of the four positions within two nucleotides upstream or downstream of the target Ψ site, and evaluated the enzymatic reaction efficiency on these oligonucleotides using MALDI-TOF MS. In agreement with the ELAP-seq result, Mj1640 does not possess obvious selectivity for nucleobases at -2, -1 and +2 positions but exhibits selectivity for nucleobase at the +1 position, where uridine is preferred over other nucleobases (Figure S4). These results are consistent with the previous report that Mj1640 requires 5’-Ψ (U/ Ψ)-3’context for optimal catalysis^27^. Previous reports also suggest that a significant proportion of Ψ sites in mRNA appear in the 5’-ΨU-3’ context^18, 19^, the preferred sequence context of Mj1640, which is supported by the observation that 30-60% of Ψ in cellular mRNA could be labeled by the enzyme (Figure 1e and Figure S2). Despite of this sequence preference, our enrichment protocol allows pulling down oligonucleotides of almost all sequence contexts to various extents. Meanwhile, two independent pull-down experiments of the standard oligos resulted in highly consistent results (Linear correlation with R^2^=0.98), demonstrating excellent reproducibility of the pull-down protocol (Figure 2d).

To detect Ψ at single base resolution, we found that at 500x lower dNTP concentration, superscript III or M-MLV could be mostly arrested by the alkynylated Ψ site in synthetic RNAs with or without the addition of a biotin molecule during reverse transcription (RT) (Figure 2e). The specific enrichment of RNA fragments containing Ψ in various sequence contexts and the RT stop signature at the labeled sites together would enable ELAP-seq to identify Ψ modification sites at single base resolution.

### Validating ELAP-seq by detecting known Ψ modifications in rRNA

We first evaluated the ability of ELAP-seq to detect known Ψ modifications in rRNA^32^. We assessed two conditions in reverse transcription when constructing Illumina sequencing libraries: 1) using superscript III with 500x lower dNTP, which facilitates generation of stop signatures as shown in Figure 1e; and 2) using superscript IV with normal concentration of dNTP. As expected, the reverse transcription mediated by superscript III at 500x lower dNTP concentration was substantially arrested after putting an adenosine opposite the Ψ, generating a drastic stop signature right at the site of the modification (Figure 3a). Interestingly, superscript IV with normal dNTP concentration also generated stop signatures at Ψ, although to lesser extent, suggesting that the on-bead RT reaction used in our library construction protocol further facilitated the generation of the stop signature.

**Figure 3.**
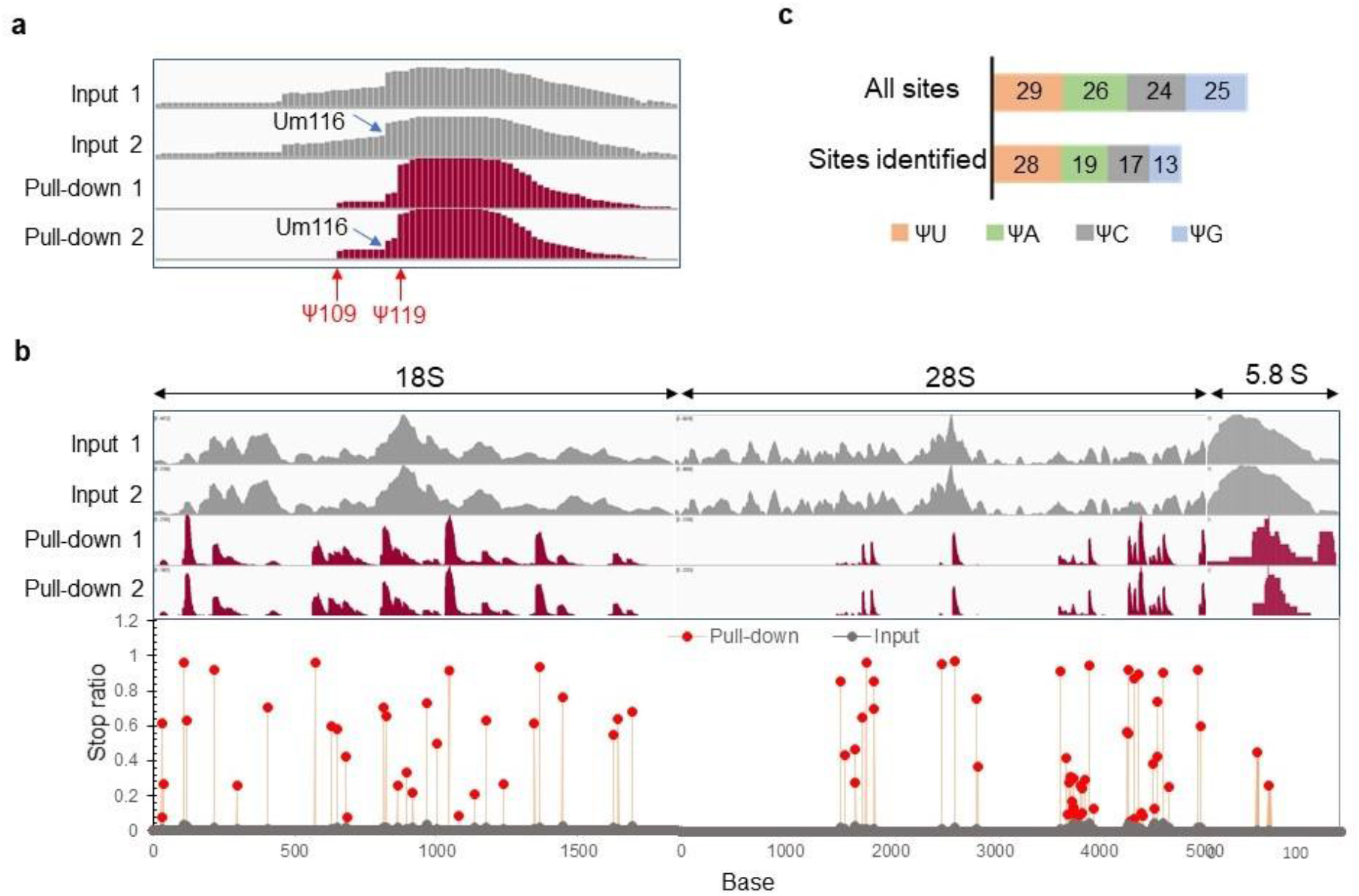
ELAP-seq detects known Ψ modification sites on rRNA. **a**) IGV snapshot shows distinct RT stop patterns between Ψ and N_m_. Stop signatures at Ψ are generated only after enzymatic treatment whereas N_m_ signatures exist in both the input and the pull-down samples. **b**) IGV snapshot showing the specific pull-down of RNA segments containing Ψ (top) and RT stop sites indicating Ψ modifications on rRNA (bottom). **c**) A summarization of identified rRNA sites in different sequence contexts.

We focused on data generated with superscript III for rRNA detection due to the higher stop ratios therefore higher resolution. As expected, Nm sites in rRNA were effectively filtered away based on the same stop signature in the input library (Figure 3a and Figure S5), whereas Ψ sites were specifically identified based on the stop signatures uniquely generated after the enzymatic treatment (Figure 3a and Figure S5). A site in the ribosome RNA is defined as a candidate Ψ site when fulfilling the following requirements: 1) the stop ratio at the site is > 0.07 in the pull-down sample and < 0.05 in the input sample; 2) arrested reads at the site in the pull-down sample is 15% higher than that at the neighboring sites to eliminate “stutter effect” caused by lower processivity of the RT enzyme near and downstream of the modification sites; 3) arrested reads at the site in the pull-down sample are > 7 when stop ratio is < 0.5 and > 5 when stop ratio is >= 0.5; and 4) the stop signature in the pull-down sample appears in both replicates. Our methods successfully identified 76 out of 104 known Ψ sites in rRNA (73% sensitivity) at single base resolution, covering 28/29 sites in the 5’ΨU3’ context as well as 48/75 sites in other contexts (Figure 3b-c). Our method also led to stop signatures at U31 and U630 of 18S, which likely originated from off-target mapping of short intronic reads to rRNA genes (Figure S5), as well as U1603 and U4561 of 28S, whose originalities are unclear (95% accuracy). Note that Ψ modifications in rRNA often cluster, which explain the inability of our method to detect the rest sites which have low stop signature resolution.

### ELAP-seq identifies ten thousand of candidate Ψ sites from human transcriptome

Next, we applied ELAP-seq to messenger RNAs of HeLa and HEK 293T cell lines (Figure S7a). Pull-down peaks of mRNAs are well isolated with low backgrounds, as exemplified by the pull-down maps on the mitochondria chromosome, highlighting the specific enrichment of RNAs containing Ψ (Figure 4a). Stop signatures are observed at Ψ modification sites under the enriched peaks, enabling the determination of the single base location of the Ψ modification (Figure 4b).

**Figure 4.**
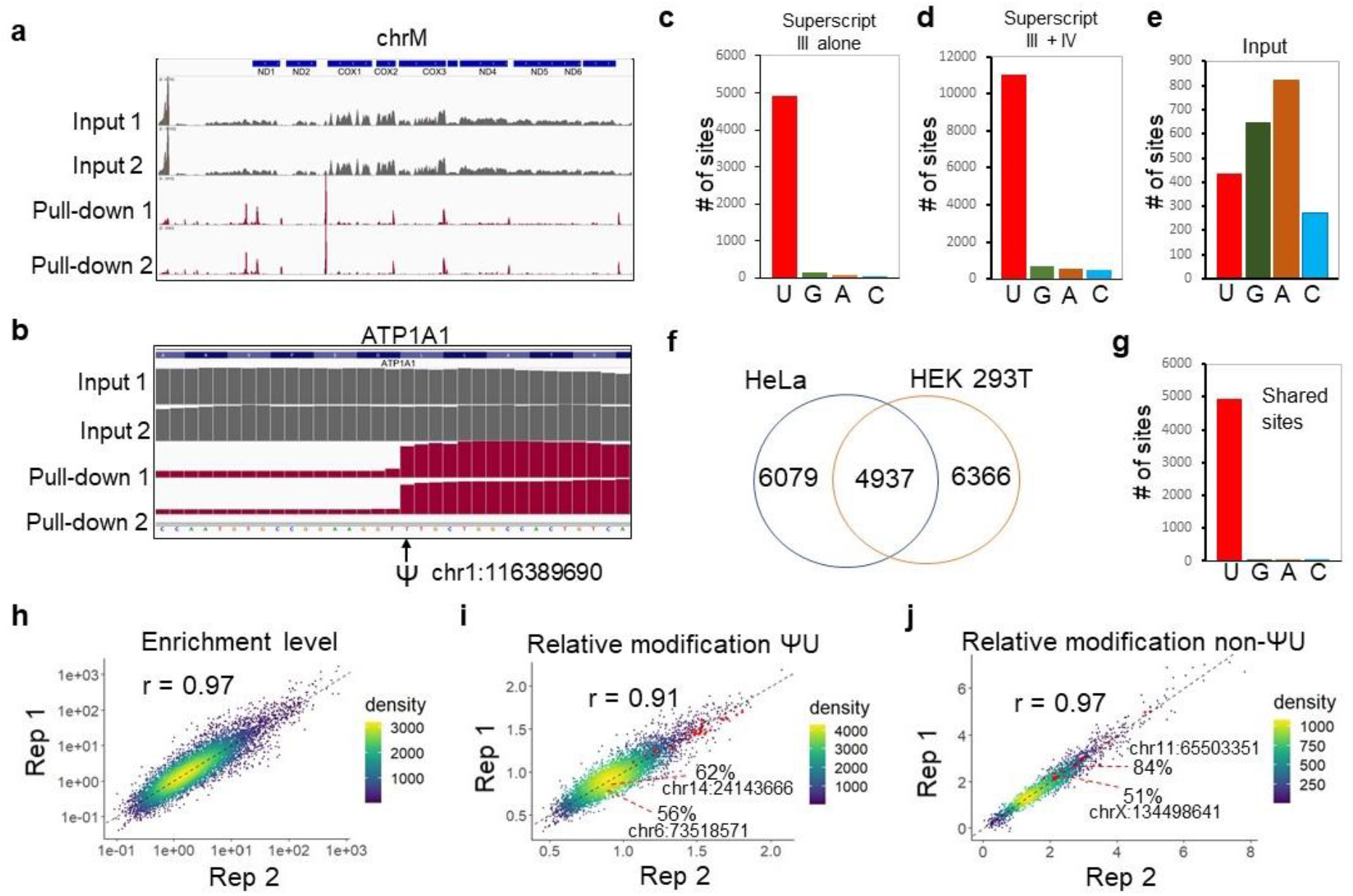
ELAP-seq identified over 10,000 candidate Ψ sites from HeLa and HEK 293T cell lines. **a**) IGV snapshot showing reads mapping to the mitochondria chromosome as an example of the pull-down profile. **b**) An example of a Ψ modification site determined based on a prominent stop signature. **c-d**) Base distribution of RT arrest sites identified from HeLa mRNAs using libraries generated with Superscript III alone (**c**) or combined reads of libraries generated with Superscript III and Superscript IV (**d**). **e**) Base distribution of RT arrest sites identified in the input samples of HeLa mRNAs. **f**) Overlap of Ψ sites identified from HeLa and HEK 293T cell lines. **g**) Base distribution of all shared sites between HeLa and HEK 293T cell lines. **h**) The enrichment levels of candidate Ψ sites in HEK293T cell line in the pull-down sample relative to the input sample are highly correlated between two biological replicates. Only sites with RPM > 2 in the input sample is used to obtain more accurate quantification. **i-j**) Estimated relative modification extent for sites in the ΨU contexts (**i**) or the non-ΨU contexts (**j**). Sites whose absolute modification levels were previously determined by CLAP method are explicitly labeled^33^.

We used an optimized pipeline to analyze the ELAP-seq data of messenger RNA, which primarily relies on parameters of stop ratio and reads coverage to define candidate modification sites (See Figure S6 and the Method section). Importantly, we did not include any steps that may biasedly select for a nucleoside over others, thus the percentage of U sites among all identified sites can be used as an indicator of rate of false positives. To achieve a balance between the number of sites identified and the rate of false positives, we optimized cutoffs for stop ratio and reads coverage and allow for gradients of coverage for different ranges of stop ratios. With our pipeline, 5,195 sites were identified from HeLa cells using data from libraries built with superscript III, which are composed of 4930 U (94.8%), 144 G (2.7 %), 75 A (1.4 %) and 46 C (0.88%). The predominant enrichment of U validated the specific identification of Ψ using our protocol (an estimated accuracy of 98%) (Figure 4c). We noticed that some sites that were covered with insufficient reads in one of two replicates were filtered off although they share highly similar stop pattern with the other replicate; simply lowering the cutoff for reads coverage to decrease the rate of false negatives would also increase the level of false positives. By comparison, superscript IV leads to higher reads coverage yet lower stop ratios than superscript III. With combined reads of superscript III and IV which gave better signals, 7,305 more sites were identified and together we identified 12,677 sites from HeLa cell line, which are composed of 11,016 U (86.8%), 660 G (5.2%), 553 A (4.3%) and 448 C (3.5 %) (an estimated accuracy of 95%) (Figure 4d). To eliminate the possibility that uridine is biased by factors other than Ψ modification, we also analyzed input samples using the same pipeline and no enrichment for U was observed (Figure 4e).

The reads per million (RPM) coverage of identified sites in input samples ranges from 0.09 to 791, demonstrating the capacity of this method to identify Ψ modifications on genes with a wide range of expression levels (Figure 4h). We set out to validate several candidate Ψ sites using CMC based RT-qPCR method. Among the 13 sites that produced reliable qPCR signals, 10 were successfully validated (Figure S7b-c). The ones not successfully validated probably have low levels of modification or labeling that is below the detection limit of CMC RT-qPCR method without enrichment.

We performed the same analysis for HEK 293T cell lines and identified 11,303 candidate Ψ sites, among which 4,954 sites were also found in HeLa cell line, accounting for 97% of all shared sites between the two cell lines (Figure 4f-g and Figure S7d), indicating that they are highly confident sites that are conserved between these two cell lines. Note that some sites passing the filter in one cell line but found in only one replicate in the other cell line were filtered off. Therefore, the reported number of Ψ modification in this study may still underestimate the true number of modifications. More replicates would effectively increase the number of sites identified.

Sites identified in our study overlapped with 40-42%, 49%, 27%, 17% and 12% of sites identified by BID-seq, RBS-seq, CeU-seq, Ψ-seq and PseudoU-seq, respectively (Figure S8)^16–19, 21^. The higher overlap with bisulfite treatment-based methods (i.e., BID-seq and RBS-seq) over CMC-based methods indicates that the former strategies outperform the later ones to identify true positive sites.

To assess the capability of the method to potentially quantify the extent of modification, we first calculated the enrichment level of each site by dividing normalized read coverage RPM at the modification site in the pull-down sample by that in the input sample to account for the gene expression difference. This parameter showed Pearson correlations of higher than 0.95, demonstrating an excellent reproducibility of the method (Figure 4h and Figure S9a) and the feasibility to use this parameter for differential modification comparisons across different conditions. We further looked for intra-group quantification. We reasoned that the enrichment level is determined by the combined effects of the modification level and the enzyme’s sequence preference. By performing pull-down and qPCR on a synthetic oligonucleotide modified to various levels, we found an exponential correlation between the modification level and the enrichment level, based on which a fit equation was deduced to determine the relative modification level of each site taking into consideration the enzyme’s sequence preference (Figure S9b).

Due to the large discrepancy in enzymatic preference between the 5’ ΨU 3’ context and other sequence contexts, we categorized the sites into ΨU and non-ΨU groups. Supporting the semi-quantification capability of the method, Ψ sites in rRNAs, which are known to be modified to nearly 100%^32^, are distributed at the higher end of the relative modification level spectrum in each group (Figure 4i-j). We also correlated the parameter of relative modification level of several mRNA modification sites to the absolute modification levels previously quantified by CLAP, and found a positive correlation (Figure 4i-j)^33^, indicating the method may distinguish highly modified sites from lowly modified ones. Based on this, our data suggests that a considerable proportion of sites (∼1/3) identified here are modified to levels higher than 50%. Note that the quantification may not be valid for clustered sites; RT arrests would skew the actual enrichment level, leading to weaker correlation between enrichment levels and the modification levels.

### New insights into Ψ biosynthesis and function

The 11,016 candidate Ψ sites identified from HeLa are distributed in 3,856 transcripts, of which 73% (8,041 sites) are located in messenger RNAs (Figure 5a), with enrichment in CDS and 3’ UTR and depletion in 5’ UTR, which agrees with previous studies (Figure 5b)^16^. Most modified transcripts (2449) harbor only one Ψ, although some transcripts are heavily modified including *FLNA* (42 sites), *A2M* (35 sites), *ATP1A1* (26 sites), *GANAB* (25 sites) and *PRKDC* (21 sites) (Figure 5c). By analyzing the sequence motif of 10 nucleotides surrounding the candidate Ψ sites, we found that the identified sites are enriched in the 5’ΨU3’ context, which reflects the sequence preference of the enzymatic labeling (Figure S10a). Interestingly, the motif exhibits a preference for uridine on the 5’ side, which is not the intrinsic preference of the enzyme and may reflect the preferred sequence motif of the Ψ installation machinery (Figure S10a-b). Importantly, we found that Ψ exhibits higher trend to locate in mismatched and UG or GU wobble paired positions compared to unmodified U sites (Figure 9d and Figure S10c), which is consistent with the previous report that Ψ installed by PUS7 tends to occur in less structured RNA^34^. This observation is unlikely due to a structural preference of the enzymatic labeling because: 1) we used a protocol that disrupts RNA secondary structures prior to enzymatic labeling; 2) sites in matched positions in rRNA were as effectively detected as sites in mismatched and wobble positions (Figure S10d); and 3) the enzymatic labeling efficiency on isolated total RNA at 37 °C is the same as that under the temperature of 65 °C (Figure S10e), although the enzyme activity is higher at 65 °C.

**Figure 5.**
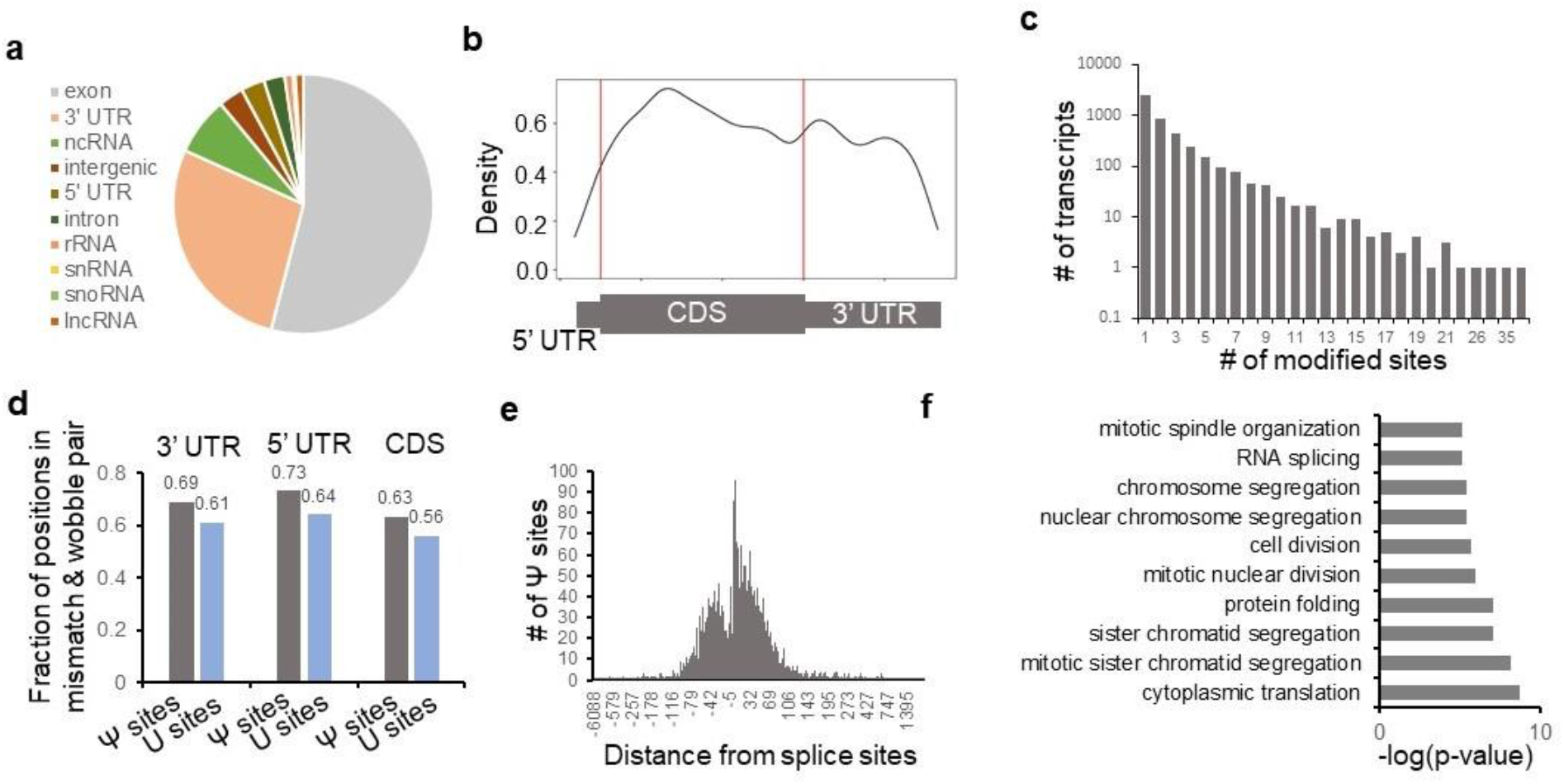
New insights into Ψ distribution and function. **a**) Distribution of candidate Ψ sites identified from HeLa cells in different genomic regions. **b**) The metagene plot of candidate Ψ sites identified from HeLa cells along the mRNA gene. **c**) Numbers of Ψ modification sites harbored by modified transcripts. **d**) Ψ modification prefers to locate in less-structured regions compared to unmodified uridine. **e**) Distance of Ψ sites from the nearest splice sites. **f**) Top 10 clusters of GO terms of biological process for highly modified RNA transcripts.

Additionally, we observed Ψ locates within 100 nucleotides to the splicing sites, suggesting a correlation between the modification and splicing (Figure 5e), as previously reported^5^. Gene ontology analysis shows that the top 10 clusters of molecular functions or biological processes of highly modified mRNA transcripts (with estimated modification levels > 50%) include mitotic cell cycle transition, protein post-translational modification, protein localization, RNA binding and translation (Figure 5f and S10f). Further studies are required to more thoroughly interrogate the determinants of the biosynthesis of Ψ and its biological function.

## Discussion

There are limited tools to study Ψ modifications and our understanding of functions of Ψ are still limited. Here we introduce a methyltransferase-mediated labeling strategy that specifically tag Ψ with chemical groups in a one-step mild enzymatic reaction. Although the enzymatic reaction catalyzed by the N1-methyltransferase Mj1640 has been reported to only occur at Ψ54 of tRNA in *Methanocaldococcus jannaschii*^26^, our biochemical characterization showed that the enzyme is actually promiscuous *in vitro* without stringent requirement for RNA secondary structures or specific sequences. It displays excellent specificity towards Ψ over U, and importantly, it readily adopts propargyl-SeAm as a cofactor to install a propargyl group onto Ψ at position N1, allowing further manipulations through clicking on various functional modules.

Based on these results, we introduced a robust method named ELAP-seq, which enriches RNA fragments bearing Ψ for detection at single base resolution transcriptome wide. We found that the enzyme prefers to catalyze methylation of Ψ that has a U at adjacent 3’ position, but Ψ in other sequence contexts can also be labeled to various extents, enabling detection of Ψ in almost all sequence contexts. The difference originated from sequence preference can be normalized using data of standard oligonucleotides bearing the specific sequence contexts for semi-quantitative comparisons of inter- and intra-group modification variations. This method is conceptually distinct from all previous methods that rely on relatively harsh chemical treatments. ELAP-seq labels and enriches Ψ modified RNA fragments under physiological conditions for sequencing using relatively low amount of starting material with low sequencing cost. These advantages allowed reliable identification of over ten thousand candidate Ψ sites in transcripts in HeLa and HEK 293T cells with low levels of noise signals. The high number of identified sites are not unexpected, as the enrichment allows detection of lowly modified sites and LC-MS/MS data already suggested a similar abundance of Ψ to that of *N*^6^-methyladenosine (m^6^A) in mRNA, another highly prevalent modification found at thousands of sites in the transcriptome^35–37^.

The sites identified by ELAP-seq provided important insights into Ψ deposition and function. Our findings show that Ψ tends to locate in less structured regions. This may attribute to the preference of certain Ψ synthases, such as PUS7 catalyzing Ψ formation in less structured regions^34^. The gene ontology terms of highly modified sites suggests a close correlation of Ψ to translation. This has been supported by previous observations that incorporations of Ψ in mRNAs and tRNAs regulate translation in a context-dependent manner^4, 38, 39^. Considering that Ψ is widely thought to be used by cells to increase their adaptivity to environmental challenges^40, 41^, our observations suggest a role for Ψ in providing cells with a mechanism to rapidly alter protein synthesis in response to environmental stimulations.

Pseudouridylation is irreversible and there has not been a good way to interrogate functions of specific Ψ sites inside cells. The activity of the overexpressed Mj1640 enzyme in cells suggest that this enzyme could be fused to inactive dCAS13 and employed for site-specific conversion of Ψ to m^1^Ψ at endogenous sites. This methodology may provide a promising avenue to connect specific Ψ to biological function and modulate functions of Ψ-modified transcripts if the N1 position is critical to the native role of specific Ψ. Additionally, the enzymatic treatment may contribute to nanopore sequencing by further decorating Ψ with bulky functional groups and allowing enrichment of Ψ-containing transcripts. Looking forward, we anticipate more to-be-discovered enzymatic labeling approaches including ours to advance basic research as well as the development of new therapeutic strategies to advance biomedicine in the future.

## Materials and Methods

### Protein purification

The Mj1640 plasmid for overexpression in E. coli was a kind gift of the Jens Wöhnert laboratory at the Goethe-University Frankfurt. In the plasmid, Mj1640 gene with an N-terminal His-tag optimized for overexpression in *E. coli* was cloned into the pET11a overexpression vector. The plasmid was expressed in T7 express competent E. coli cells (NEB) grown in LB medium and overexpression of the protein was induced by adding IPTG to 1 mM at OD6_00_ of around 1 and incubating for 3 h at 37 °C. Two liters of induced cell cultures were pelleted and resuspended in 60 mL lysis buffer composed of 50 mM sodium phosphate (pH 7.5), 500 mM NaCl, 10 mM imidazole, 5 mM beta-mercaptoethanol, and 1 tablet of complete protease inhibitor (Roche). Cells were lysed in French Press with 1-2 rounds of lysis under the pressure between 15 000 psi and 20 000 psi. The purified lysate was run through a Ni-NTA column manually, washed with 300 mL wash buffer containing 2.5 mM sodium phosphate (pH 7.5), 12.5 mM NaCl and 30 mM imidazole, and eluted sequentially with three elution buffers containing 25 mM sodium phosphate (pH 7.5), 125 mM NaCl, and 100 mM, 200 mM and 300 mM imidazole. To thoroughly remove bound nucleic acids and contaminating proteins, the eluted protein was further purified sequentially with the anion exchange column twice and Heparin column once. Specifically, proteins were diluted with buffer A (20 mM Tris-HCl, pH 7.4) to lower NaCl concentration to below 75 mM and then loaded onto HPLC-mono Q or -Heparin column. After washing with buffer A, the proteins were eluted with a gradient mixture of buffer A and buffer B (20 mM Tris-HCl pH 7.4 and 1M (for the Mono Q column) or 1.5 M (for the heparin column) NaCl). The proteins eventually eluted from the Heparin column were buffer exchanged to the storage buffer (20 mM Na_2_HPO_4_ PH 7.5, 150 mM KCl), concentrated to 20 µL, flash frozen in liquid nitrogen and stored at -70 °C. The resulting protein was > 95% pure.

### In vitro methylation reaction

The condition used for testing the reactivity on synthetic RNA oligos is 15 mM sodium phosphate (pH 7.5), 100 mM KCl, 500 µM SAM or pro-SeAm, 200 µM RNA substrate and 220 µM enzyme in a 50 µL scale reaction for various time periods at 65 °C. Synthetic RNAs were purified via phenol-chloroform extraction and ethanol precipitation. The condition used for labeling isolated cellular RNA for sequencing library preparation is15 mM sodium phosphate (pH 7.5), 100 mM KCl, 321 µM pro-SeAm (1.5 mM of Pro-SeAm dissolved in water containing 0.5% TFA (trifluoric acid), with pH adjusted using Tris-HCl pH 8.3 by mixing 20 µl Pro-SeAm, 20 µl H_2_O and 8 µl 1M Tris-HCl pH 8.3), 12.5% PEG8000, 0.8 U/µl RNaseOut, 0.8 U/µl SUPERase•In™ RNase inhibitor, around 200 ng mRNA (around 70 µM for 200 nt average size) and 64 µM enzyme in 40 µL scale reaction for 2 h at 37 °C. Cellular RNAs were purified using RNA Clean and Concentrator kit (Zymo Research) and stored in RNase-free H_2_O.

### MALDI-TOF

Purified RNA oligonucleotide from enzymatic labeling reaction was resuspended in 5 μL water, mixed with 40 μL ion exchange resin (Bio-RAD), and left at room temperature for 30 min to remove salts. Loaded 1 μL of supernatant of the mixture to a MALDI sample plate, mixed with 1 μL THAP matrix freshly added with salt, and pipet to mix until crystal is formed. The MALDI TOF MS recorded the m/z signals using negative reflector mode.

### Mammalian cell culture and RNA preparations

HEK 293T and HeLa cell lines were grown and maintained in DMEM supplemented with 10% FBS and 1% pen-strep antibiotics under 37 °C with 5% CO_2_. RNAs were extracted from cells using Trizol reagent and polyA RNA were extracted using Dynabeads® mRNA DIRECT™ Kit for two rounds. Messenger RNAs were extracted with an additional ribosome RNA removal step using the RiboMinus™ Eukaryote Kit v2. Only cells within passage number of 10 were used for all experiments.

### LC-MS/MS

RNA was digested with 3 μl 10× Reaction Buffer (100 mM NH^4^OAc, pH = 5.3) and 1 μl Nuclease P1 (>1U/ μl) in 25 μl solution at 42 °C for 2 h to beak the phosphodiester bond, after which phosphate was removed by adding1 μl FastAP (1U/μl) (ThermoFisher, EF0654) and 3 μl 1M NH_4_HCO_3_ (pH unadjusted), and incubating at 37 °C for 2 h. Upon the completion of incubation, diluted the reaction mixture to 50 μL with water and filtered the samples through 0.22 μm Millex-GV PVDF filters (Millipore). Five microliter sample was injected into ZORBAX SB-Aq 4.6 x 50 mm column (Agilent) on UHPLC (Agilent) coupled to a SCIEX 6500+ Triple Quadrupole Mass Spectrometer in a positive electrospray ionization mode. The nucleosides were quantified based on the nucleoside to base transitions: 268.0 to 136.0 (A), 284.0 to 152.1 (U), 245.1 to 125.0 (Ψ), 259.1 to 139.0 (m^1^Ψ) and 283.1 to 163.0 (p^1^Ψ). At least two independent biological replicates were used for Ψ level quantification, and each sample was injected 2 times.

## ELAP-seq

### RNA fragmentation, enzymatic labeling, end repair and click reaction

RNA was first fragmented using 10x fragmentation buffer (thermos fisher) in a 10 µL scale reaction containing 1x fragmentation buffer and 200-1000 ng RNA. The fragmentation reaction was incubated at 70 °C for 12 min to fragment the RNA to 60-200 nt. RNA was recovered using Oligo Clean and Concentrator kit (Zymo Research) and eluted in water. The recovered RNA was denatured at 90 °C for 3 mins and quickly set on ice to break RNA secondary structures. Aliquot 1/10 RNA as input.

The remaining RNA was then labeled in *in vitro* methylation reaction using pro-SeAm as co-factor following the protocol mentioned above. RNA was purified using RNA Clean and Concentrator kit, eluted in 9 µL of water, and entirely used in a 50 µL scale end repair reaction containing 1x T4 PNK buffer (NEB B0201S) and 5 µL T4 PNK (Thermos Fisher) for incubation at 37 °C for 1h. The RNA was then purified using Oligo Clean and Concentrator (Zymo Research) and eluted in 15 µL water.

Used half of the samples in a 100 µL scale click reaction containing 250 µM biotin-azide, 0.4 mM CuSO_4_, 4mM THPTA and 4 mM sodium ascorbate which is added the last. Used the other half in a control reaction where THPTA and CuSO_4_ were omitted. Incubated the reactions at room temperature with shaking at 600 rpm for 1.5 h. RNA was recovered by ethanol precipitation at -80 for overnight. The input RNA was treated with end repair reaction and used directly in RNA ligation.

### RNA ligation, pull-down and reverse transcription

RNA recovered from ethanol precipitation was resuspended in 9 µL water and mixed with 1 µL of 30 µM 3’ RNA linker (5′rApp-NNNNNAGATCGGAAGAGCGTCGTG-3SpC3). The mixture was denatured at 72 °C for 2 mins and then quickly set on ice. Added to the mixture T4 ligation buffer to 1X, PEG8000 to 15%, 0.8 U RNAsin, and water to a total of 23 µL. Mixed thoroughly, added two µL T4 RNA ligase 2 truncated KQ and mixed again. Incubated the mixture at 25 °C for 2 h and then 16 °C for 12 h.

Aliquoted MyOne C1 magnetic beads suspension (5 µL per sample) and washed the beads following the manual. Blocked the beads with 2 volumes of 1x W&B buffer containing 0.5 mg/ml BSA and 50 µg/ml tRNA for 1 h at 4 °C. Resuspended the beads in 2 volumes of pull-down buffer containing 10 mM Tris-HCl (pH 7.5), 2 M NaCl, 1 mM EDTA, 8.8 µM biotin-azide, 44 µg/mL yeast tRNA, 0.44 mg/mL BSA and 0.1% Tween-20. When working with beads, pay attention when rinsing and pipetting beads to minimize loss of beads due to sticking to the tube wall or tips.

Mixed 17 µL ligation product with 10 µL bead suspension, and added 15 µL of 0.73x IP buffer (without biotin-azide) to decrease NaCl concentration to 1 M to facilitate biotin-streptavidin binding and decrease PEG concentration to below 5% to avoid bead aggregation. Incubated the mixture for 1 h at 4°C and then 30 mins at room temperature. Washed the beads 6 times with 100 µL 1x W&B buffer supplemented with 0.05% Tween-20, and then 2 times with 50 µL 1x W&B buffer with each wash incubated at 4°C for 5 mins. Changed tubes every three washes. Changed tubes again after the final wash and resuspended beads in 12 µL 10 mM Tris-HCl.

For input RNA samples, dilute each ligation reaction to 47 µL with water. To remove un-ligated adapters, first added 2 µL 5’ deadenylase and incubated at 30 °C for 1 h, then added 1 µL RecJ and incubated at 37 °C for 1 h. Purified the RNA using RNA Clean and Concentrator kit (Zymo Research) and eluted the product in 12 µL water.

Six µL beads from pull-down or 6 µL purified RNA from input was used in reverse transcription (RT). Set up RT reaction by adding 1 µL 2 mM RT primer (5′-ACACGACGCTCTTCCGATCT-3′), 1 µL 20 µM dNTP and water to 13 µL. For Superscript IV, used 10 mM dNTP instead. Denatured the mixture in a shaking incubator at 65 °C for 5 min with shaking at 900 rpm and then quickly set on ice for at least 1 min. Added 4 µL 5x SSIV buffer (for the Superscript IV) or 5x first strand synthesis buffer (for the Superscript III), 1 µL 100 mM DTT, 1 µL RNaseOut, and 1 µL Superscript IV or Superscript III reverse transcriptase, and incubated at 50-55 °C for 10 min (for Superscript IV) or 50 min (for Superscript III) with shaking at 900 rpm. After the RT reactions, washed beads once with 100 µL high salt buffer composed of 20 mM Tris-HCl pH 7.4, 1 M NaCl and 0.1% tween-20, and once with 50 µL low salt buffer composed of 20 mM Tris-HCl pH 7.4, 100 mM NaCl and 0.1% tween-20. Resuspend beads in 18 µL buffer composed of 50 mM Tris-HCl pH 8.3, 75 mM KCl, 3 mM MgCl_2_ and 0.1% tween-20 and denatured the RT enzymes by heating at 80 °C for 15 mins. After cooling the reaction on ice, added 1 µL 200 mM DTT and 1 µL RNase H and digested at 37 °C for 30 min with shaking at 900 rpm. Heated the mixture at 72 °C for 2 min and quickly set on the magnet to collect the supernatant. cDNA was purified from the supernatant using Oligo Clean and Concentrator and eluted in 8 µL water.

### cDNA ligation and PCR

All cDNA was used to mix with 1x T4 RNA ligase buffer, 0.81 µM adapter (5′Phos-NNNNNAGATCGGAAGAGCACACGTCTG-3SpC3), 1 mM ATP, 24% PEG8000, 7.5% DMSO and water in a 22 µL scale reaction, and incubated at 72 °C for 2 min before quickly chilling on ice for 2 min. One µL of 50 mM Co(NH_4_)_6_Cl_3_ and 1.8 µL T4 RNA ligase 1 (high concentration) were added and the ligation reaction was allowed to proceed at 25 °C for 12 h, followed by protein denaturation at 65°C for 5 min. The ligation product was purified with 1.9X Ampure beads and eluted in 10 µL water.

Two µL ligation product was used in qPCR to estimate the number of cycles needed in PCR. The rest of ligation product was amplified in a 25 µL scale PCR reaction containing LongAmp Taq 1x mix and 0.24 mM forward and reverse primers. To minimize adapter dimers, PCR products of the pull-down samples reverse transcribed using Superscript III were first purified with 0.9X Ampure beads, further amplified for 6 cycles, and then loaded on a 3% low melting-point agarose gel to remove primer dimers. For input samples and pull-down samples built from superscript IV, PCR products were directly run on a 3% low-melting point agarose gel. Excise the bands in the desired range and perform gel extraction to recover pure DNA for Illumina sequencing PE150. 25M reads were obtained from each sample.

## ELAP-seq Data analysis

### Spike-in data analysis

Paired end reads were merged using bbmap and deduplicated. 5nt and 6nt were trimmed respectively on the 5’ and 3’ ends to remove UMI with the option of -q 10, 10 to further trim both ends with quality cutoff of 10. Reads of spike-in oligonucleotides were extracted and sorted using the command grep -o "GCTCGCAG.*A" data.fq.gz > output.txt <output.txt sort | uniq -c > output1.txt. Subtotals of number of reads for sequences with identities varying at 2 nucleotides upstream and downstream of the Ψ site were summarized. Enrichment level was determined by the dividing the RPM in the pull-down sample to the RPM in the input sample. The heatmap for enrichment levels of all 256 sequences was plotted using R.

### rRNA data analysis

Reads R2 are used for analyzing data of biological RNA. Poly-A RNA ELAP-seq data also contain rRNA reads thus was used for analyzing Ψ modifications in rRNA. Adapter sequence of “AGATCGGAAGAGCGTCGTGTAGGGAAAGAGTGT” was first trimmed from the 3’ end of poly-A RNA R2, followed by de-duplication and removing UMI of 6 nt and 5 nt on the 5’ and 3’ end respectively. By design, after trimming adapter and UMI sequences, the first nucleotide of R2 is the stop site of reverse transcription. The trimmed reads were mapped to ribosome RNA genome using hisat2 with the options of -k 1 --no-softclip --mp 1000,999 to allow only one distinct alignment, disallow soft-clipping and allow no mismatch. RT arrest was called using JACUSA2 with the options of ‘rt-arrest -m 0 -p 2 -c 1 -P FR_SECONDSTRAND’ to filter off positions with mapping quality < 0 or reads coverage < 1 and specify second strand sequenced. The resulting output file was processed sequentially with in house scripts rRNA_filter_p1.sh, Stutter_removal_rRNA.py and rRNA_filter_p2.sh to calculate stop ratios, remove false positive sites due to the stutter effect, and select for sites whose stop ratio is > 0.07 in the pull-down sample and < 0.05 in the input sample and whose arrested reads in the pull-down sample are > 7 when stop ratio is < 0.5 and > 5 when stop ratio is >= 0.5.

### mRNA data analysis

Reads R2 from samples generated with superscript III were trimmed 6 nt on the 5’ end and 5 nt on the 3’ end, whereas reads from samples generated with superscript IV was trimmed 7nt on the 5’ end and 5nt on the 3’ end as superscript IV tends to elongate for one more nucleotide. Reads after adapter trimming, deduplication and UMI trimming were mapped to hg38 using hisat2 with options of “--known-splicesite-infile --rna-strandness F --no-softclip”. To combine samples built from superscript III and IV, bam files for the same sample generated from Superscript III and IV were combined to generate a new bam file. We then used MACS2 to call highly enriched peaks and categorized all mapped sites in the pull-down samples into groups inside (in-group) and outside (out-group) of highly enriched peaks. We used JACUSA2.0 to calculate RT-arrest at each site, with parameters ‘rt-arrest -m 0 -p 2 -c 1 -P FR_SECONDSTRAND’.

We pre-processed the resulting files with the following steps: 1) For sites mapped with multiple nucleotide identities (due to opposite strand orientation and mismatches), we focus on the most abundant nucleotide; 2) we filtered away sites in any region covered by reads that all share exactly the same start and end locations, which is likely caused by off-target mapping of reads whose originality is another region at a different position in the genome containing a same sequence as the current region; and 3) we removed sites whose reads coverage were less than 1/3.5 of any nearby sites within 30 nt upstream (when reads are mapped to the - strand) or downstream (when reads are mapped to the + strand) to decrease false positives due to the effect of diminished processivity of the enzyme.

Next, we determined candidate Ψ sites based on parameters of stop ratios and reads coverage. To call as many sites as possible, at the early stage we used a low stop-ratio cutoff of 0.1 and reads coverage of 8 for all sites. We then optimized these parameters as follows: 1) we required higher reads coverage for lower levels of stop ratios to decrease false positives; 2) as sequence preference of the enzymatic labeling resulted in relatively higher reads coverage at sites in the in-group than sites in the out-group, we allowed slightly lower reads coverage for sites in the out-group. 3) using percentage of U among identified sites as an indicator, we screened cutoffs for stop ratios and coverage to find a balance between the number of sites identified and percentage of U.

We set several rules to remove noises caused by various factors: 1) to remove the noise caused by off-target mapping to repetitive regions, we only select for sites that are covered by at least one uniquely mapped read for further analysis; 2) to remove noises caused by the RT “stutter” near and downstream of the modification sites, we require that the stopped reads at a candidate Ψ site must be at least 15% higher than its neighboring sites within 6 nt upstream and downstream; after this step sites identified from superscript III alone and sites identified from combined reads of Superscript III and IV were combined.; 3) to remove noises originated from non-specific binding of RNA to the beads, we looked for consistent enrichment and stop patterns between independent biological replicates.

The obtained sites were further filtered to remove stuttered sites again and sites with average stop ratios of the two replicates higher than certain values are selected depending on whether it is inside or outside the enriched peaks and whether it is identified from superscript III sample alone or from combined reads of superscript III and superscript IV. We found that to filter stop ratios this way efficiently decrease false positive ratio. The identified sites were further filtered based on the stop signatures in the input libraries (< 0.05 in input).

Metagene analysis was performed using MetaPlotR^42^. Gene ontology enrichment analysis was done using clusterProfiler^43^.

### Validation using CMC-RT-qPCR

Total RNA was treated with Cyclohexyl-N’-(2-morpholinoethyl)carbodiimide (CMC) as previously described (10.1261/rna.072124.119). Briefly, 15 or 10 µg total RNA in 12 µl sterile H_2_O was used for the treatment (CMC +) and the control reaction (CMC-). 24 µl TEU buffer (50 mM Tris-HCl (pH 8.3), 4 mM EDTA, 7 M urea,) and 4 µl freshly prepared 1M CMC (C106402-5G, Sigma) in TEU buffer or 4 µl TEU buffer was added to the CMC+ or CMC-tube respectively. Both samples were incubated at 30 °C for 16 h to ensure complete conversion of Ψ to Ψ-CMC adduct. RNA was then purified with ethanol precipitation. The RNA pellet was then air dried and resuspended in 40 µl of 50 mM sodium carbonate and 2 mM EDTA (pH 10.4) and incubated at 37 °C for 6 h to remove other CMC adducts. RNA was purified again with ethanol precipitation, air dried and resuspended in sterile deionized H2O and stored at -80 °C before use. The RNAs were used in RT-qPCR using primers as listed in supplementary table 1.

### Estimation of modification levels

We used a synthetic oligonucleotide with sequence of 5’-GGA GAC GGU CGG GΨUC CAG AUA UUC GUA UCU GUC to study the relationship between the enrichment level and the modification percentage. Different ratios of biotin-propargyl-Ψ -modified and unmodified oligos are mixed to make a total of 0.1 pmol comprising 1%, 9%, 25%, 50% and 100% of modified oligos. Pull-down was performed followed by RT-qPCR to determine relative amount of pull-down oligos. By normalizing the enrichment level to the data of modification of 1%, an exponential correlation was found between the modification percentage and the normalized enrichment level: Y = 1.879*e^0.0469X^ where Y is the normalized enrichment level and X is the apparent modification level. Based on this equation, we propose that for a site in a certain sequence context, Y = A*e^BX^, where Y is the enrichment level of the candidate site, X is the relative modification level, and B is determined by the specific sequence context the modification is located in. To determine relative modification levels for sites in different contexts given enrichment levels, we use the pull-down data of the mixed 100% modified oligonucleotides covering 256 sequence contexts to determine parameters A and B: 1) We make the approximation that the least preference oligonucleotide in the mixture pool has an apparent modification level of 0%, which means when BX=0, Y = A. Thus “A” equals to the enrichment level of the least preference oligo (Enrichment0); 2) When X =1, B = LnY -LnA, where Y is the enrichment level of the standard oligo with the same sequence context as the studied site (EnrichmentS). Therefore, for modification site with an enrichment of Y’, Y’ = A*e^(LnY – LnA)X’^, which leads to the equation:

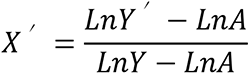

where Y’ is the enrichment level of the candidate site identified from human cell line, Y is the enrichment level of the spike-in oligo with the same sequence context as the candidate site and A is the enrichment level of the spike-in oligo with the least preferred sequence context.

### RNA structural analysis

RNA sequences of 100 nt surrounding Ψ sites with 50 nt upstream and 50 nt downstream were used for RNA secondary structure prediction using viennaRNA with the command of ‘RNAfold -p < input.fa > output.res’. Control U sites were chosen at the -100 and the +100 positions of Ψ site and RNA structures of 100 nt surrounding the control U sites were performed the same way as Ψ sites. Mfold (http://www.unafold.org/mfold/applications/rna-folding-form.php) was used to draw the secondary structures of 100 nt surrounding the chosen Ψ sites.

---

**Figure S1.**
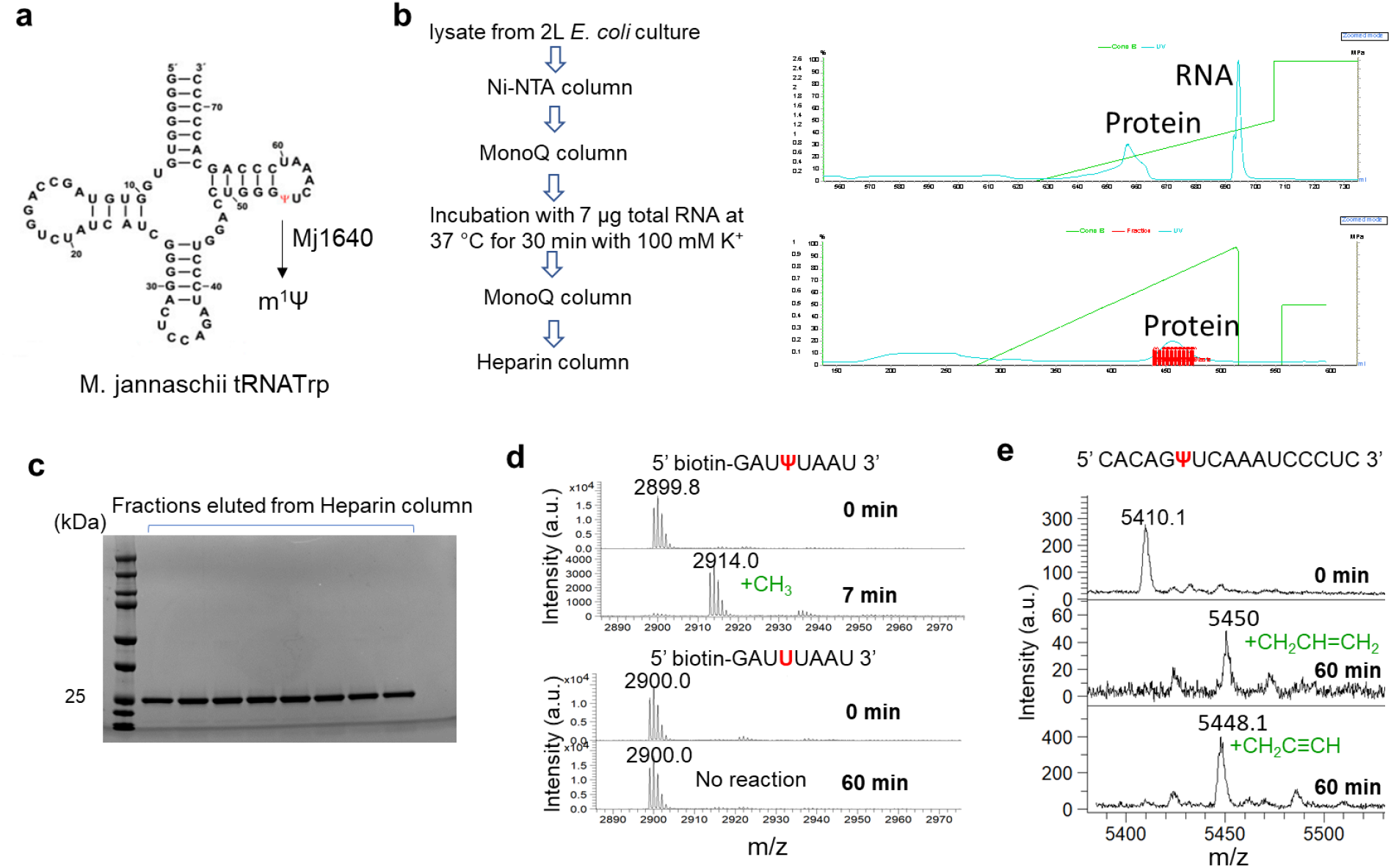
Purification of recombinant Mj1640 for *in vitro* characterization. a) Natural reaction catalyzed by Mj1640 on M. jannaschii tRNA^Trp^. b) The workflow to purify the recombinant Mj1640 (left) and HPLC trace of Mono Q chromatography and Heparin chromatography to purify Mj1640 (right). c) SDS-PAGE analysis of fractions eluted from the Heparin column, the last chromatography in the workflow. d) MALDI-TOF shows that the recombinant Mj1640 can efficiently and specifically label an 8-mer RNA oligonucleotide containing a Ψ in the center. e) MALDI-TOF shows that the recombinant Mj1640 can efficiently label Ψ-RNA oligonucleotide with an allylic or a propargyl group.

**Figure S2.**
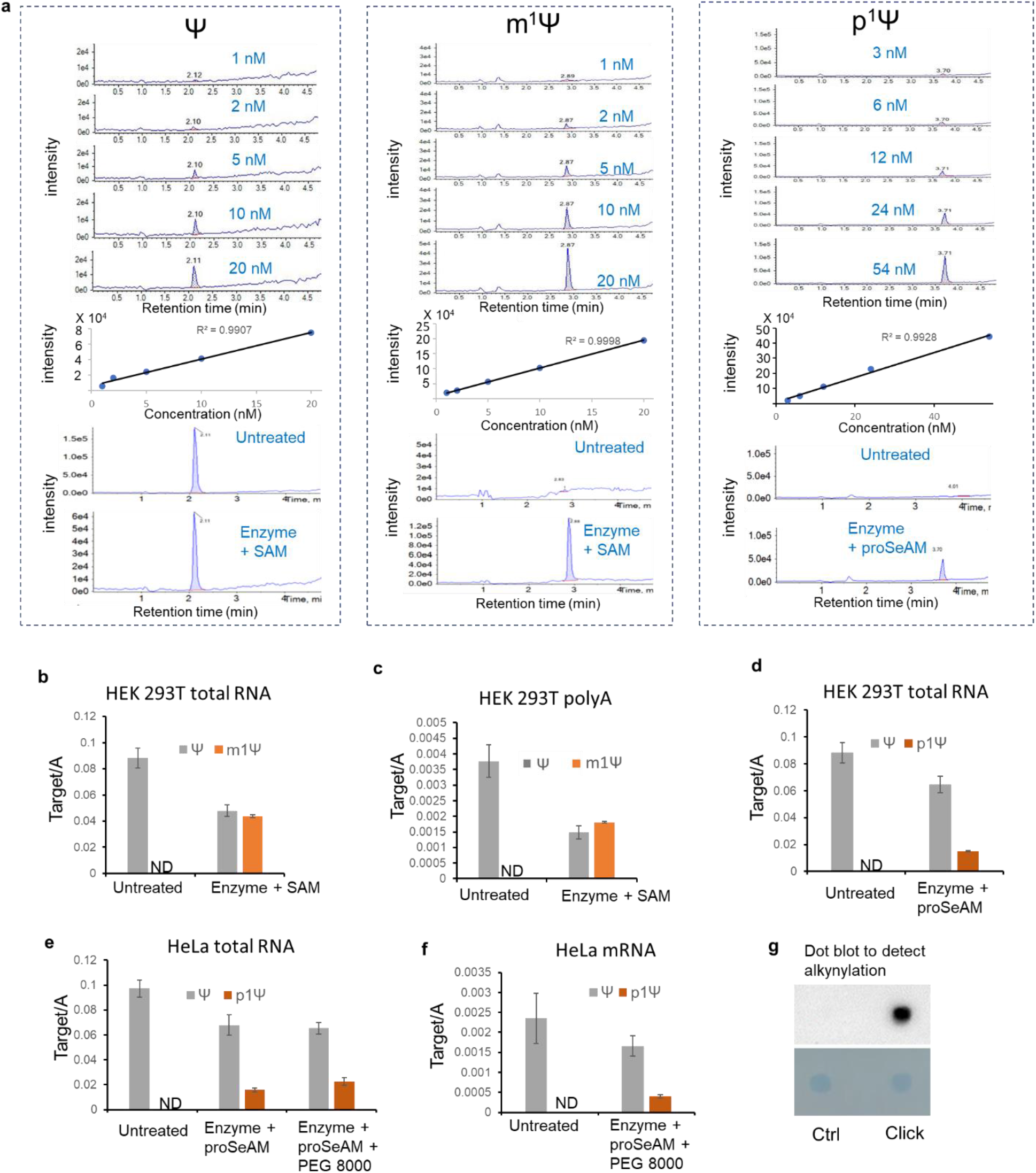
LC-MS/MS signals of Ψ, m^1^Ψ, p^1^Ψ nucleosides and quantification of enzymatic labeling of Ψ in cellular RNA isolated from HeLa and HEK 293T cell lines. a) LC-MS/MS signals and calibration curves used to quantify the efficiency of enzymatic addition on Ψ in cellular RNA with methyl and propargyl groups. b) Methylation efficiency on Ψ in total RNA extracted from HEK 293T cells. c) Methylation efficiency on Ψ in poly-A RNA extracted from HEK 293T cells. d) Propargyl group labeling efficiency on Ψ in total RNA extracted from HEK 293T cells. e) Optimization of enzymatic labeling conditions. Adding PEG8000 effectively increases the labeling efficiency. f) Propargyl group labeling efficiency on Ψ in mRNA extracted from HeLa cells. g) Dot plot method to detect alkynylation through the enzymatic labeling followed by clicking on a biotin molecule which is detected using chemiluminescent nucleic acids detection kit.

**Figure S3.**
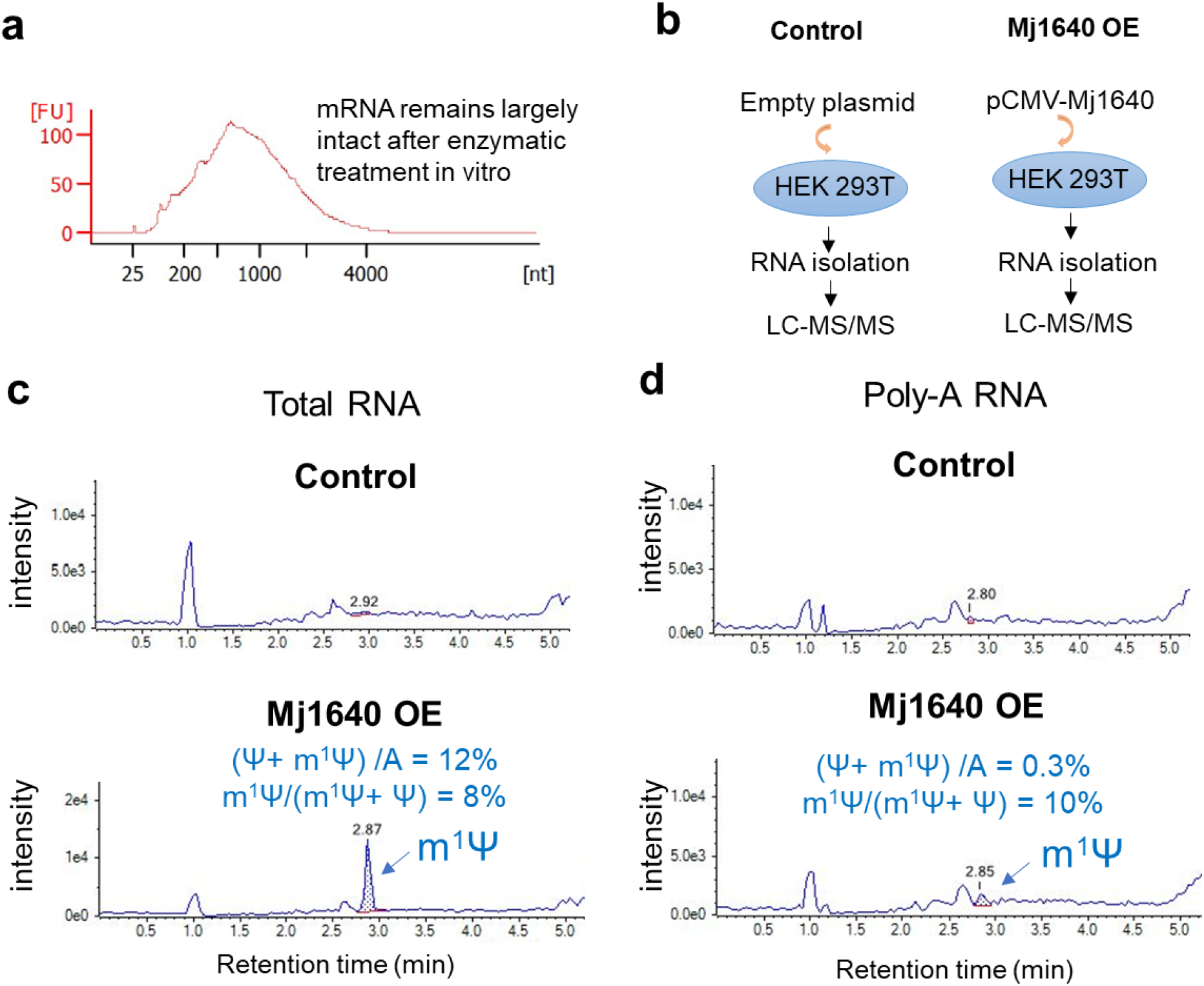
Enzymatic labeling condition with Mj1640 is mild and can also label endogenous Ψ inside cells. a) mRNA extracted from cells remains largely intact after the enzymatic treatment. Strategies to test activity of exogenous Mj1640 expressed in human cells. c) LC-MS/MS trace shows that overexpressing Mj1640 in HEK 293T cells labeled Ψ in endogenous total RNA to 8%. d) LC-MS/MS trace shows that overexpressing Mj1640 in HEK 293T cells labeled Ψ in endogenous poly A RNA to 10%.

**Figure S4.**
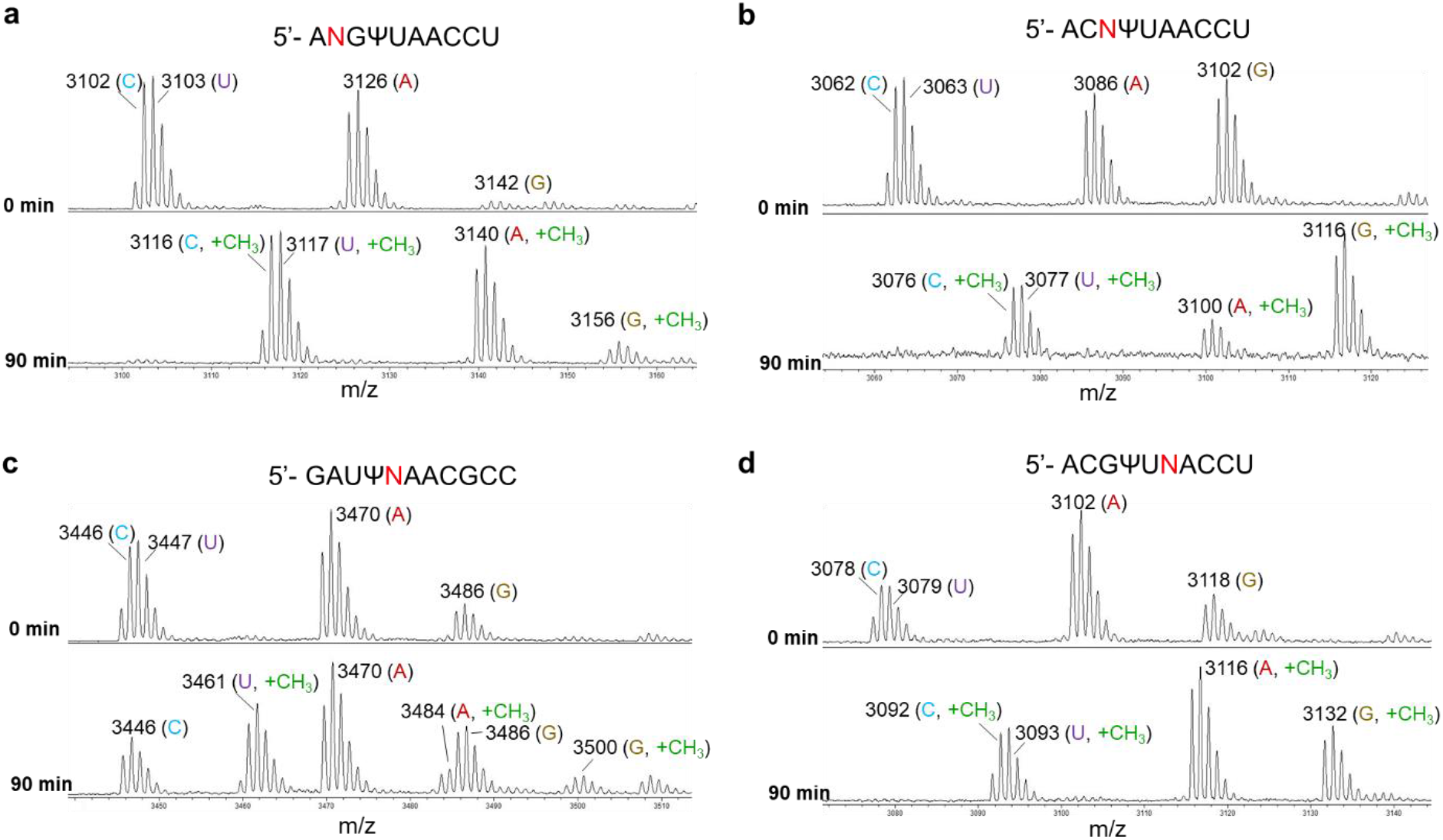
MALDI-TOF of 10- or 11-mer oligonucleotides validates Mj1640’s sequence preference for nucleobase at positions -2 (a), -1 (b), +1 (c), and +2 (d) relative to the Ψ modification.

**Figure S5.**
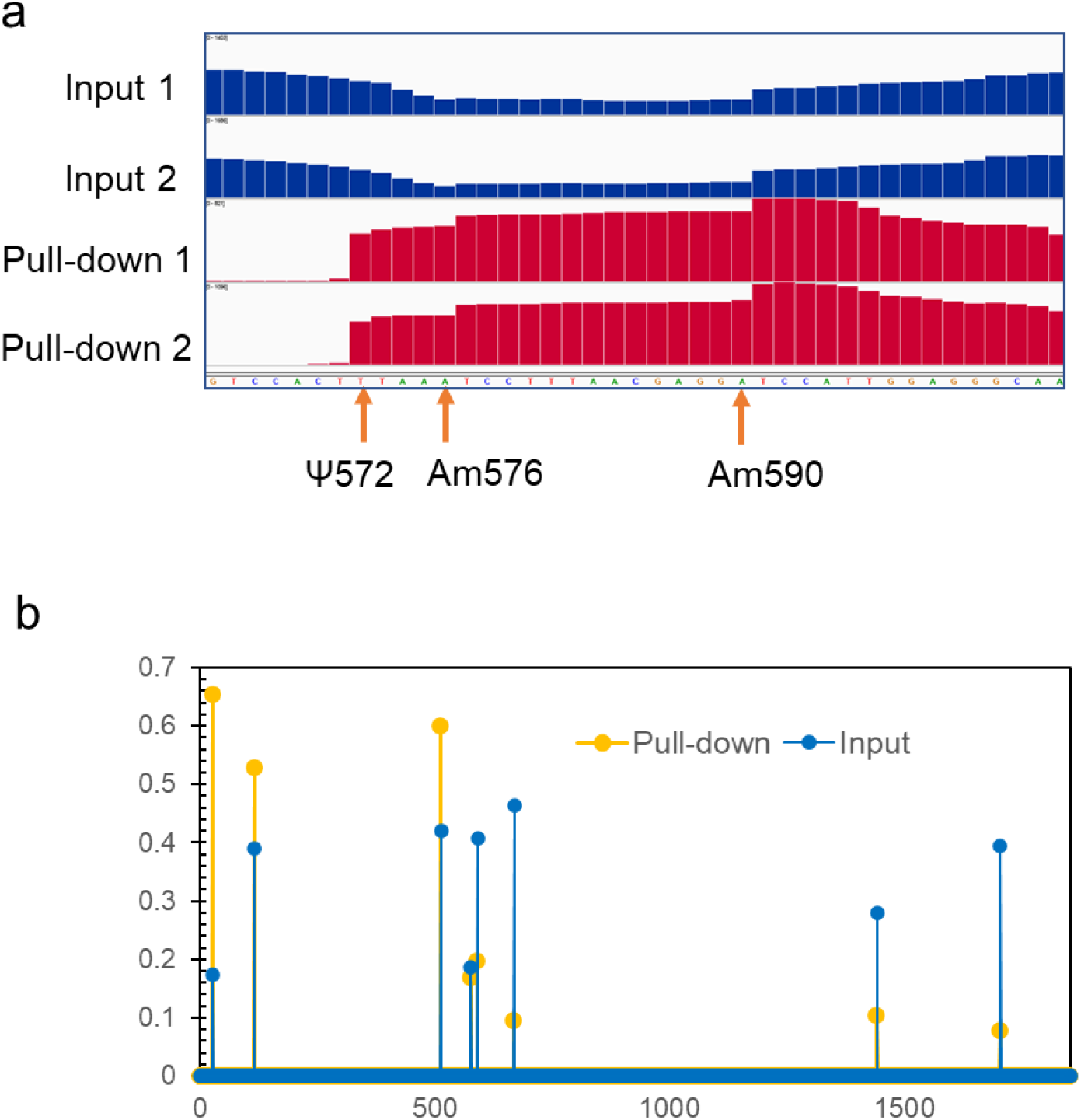
Detection of rRNA Ψ sites by ELAP-seq. a) Stop signatures at Ψ are generated only after enzymatic treatment whereas Nm signatures exists in both the input and the pull-down samples. b) Nm signatures on human 18S rRNA. The stop signatures exist in both the input and the pull-down samples thus can be distinguished from Ψ signatures.

**Figure S6.**
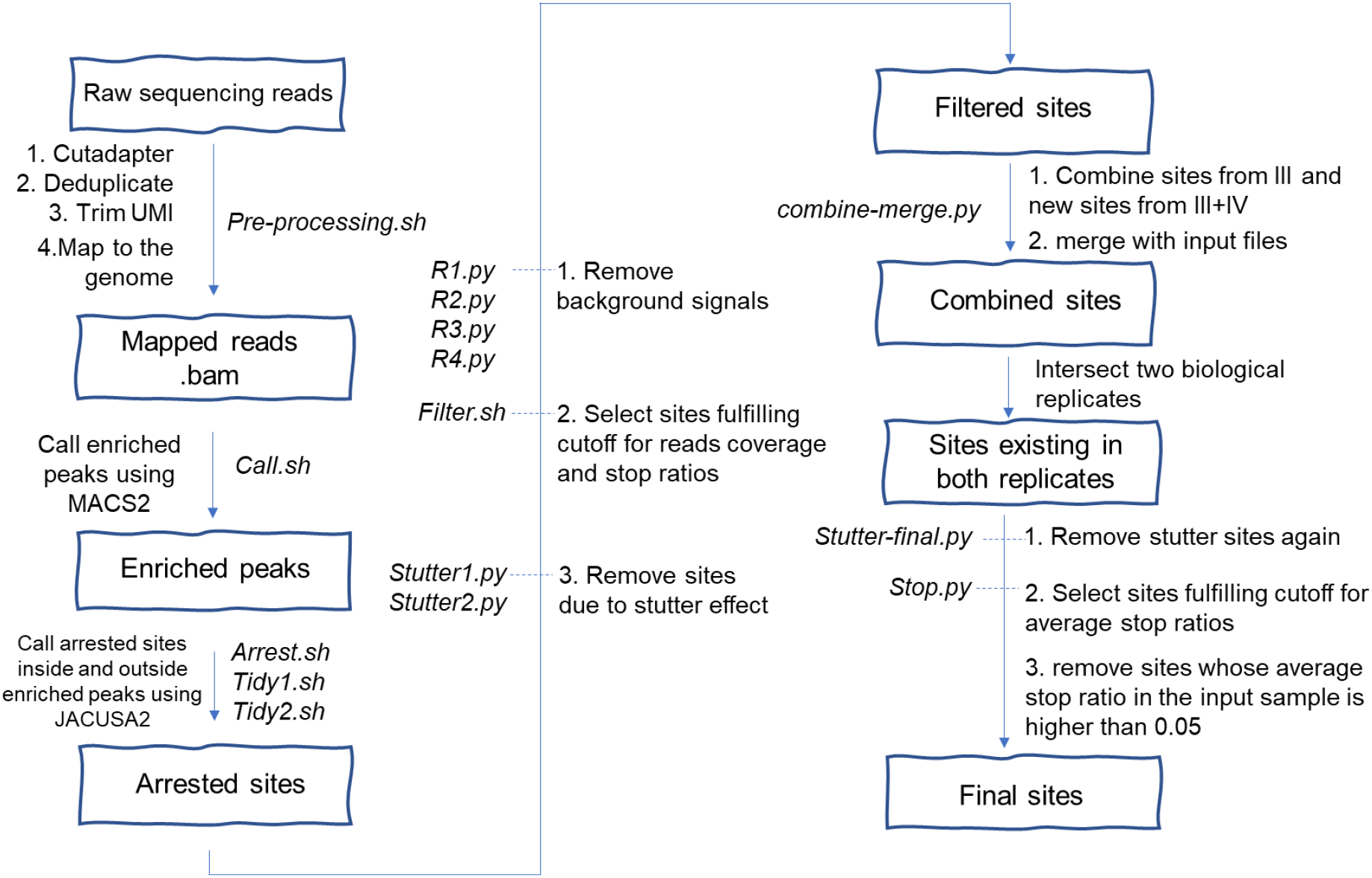
Pipelines to analyze ELAP-seq data of mRNAs.

**Figure S7.**
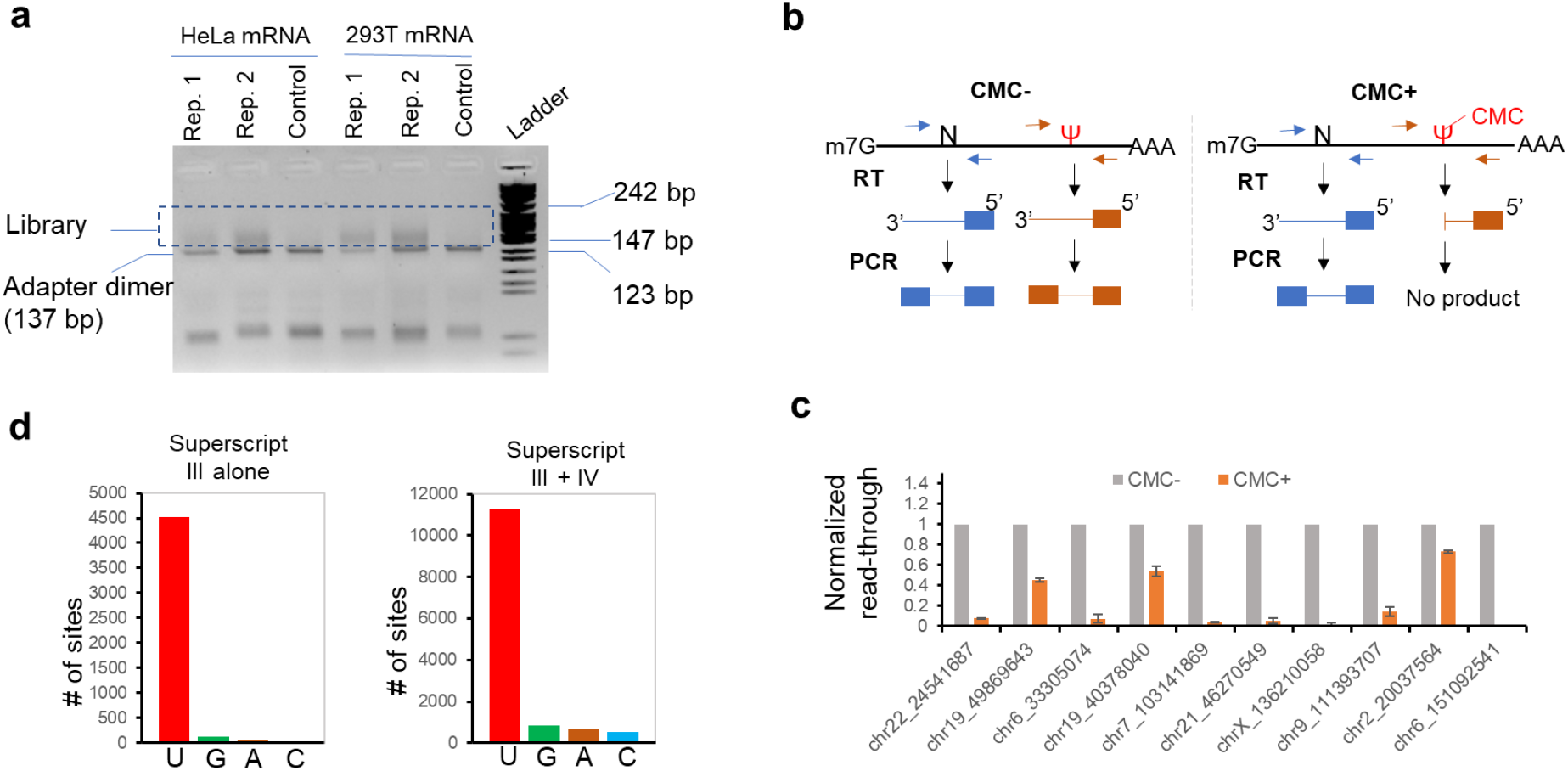
ELAP-seq identified over ten thousand candidate Ψ sites in human transcripts. a) The agarose gel of the final PCR products in the process generating libraries for Illumina sequencing. c) the scheme of using CMC-RT-qPCR to validate selected candidate Ψ sites. c) 10 candidate sites are validated by CMC-RT-qPCR. d) Base distribution of stop sites identified in mRNA extracted from HEK 293T cell lines using reads of libraries generated with Superscript III alone or combined reads of libraries generated with Superscript III and Superscript IV.

**Figure S8.**
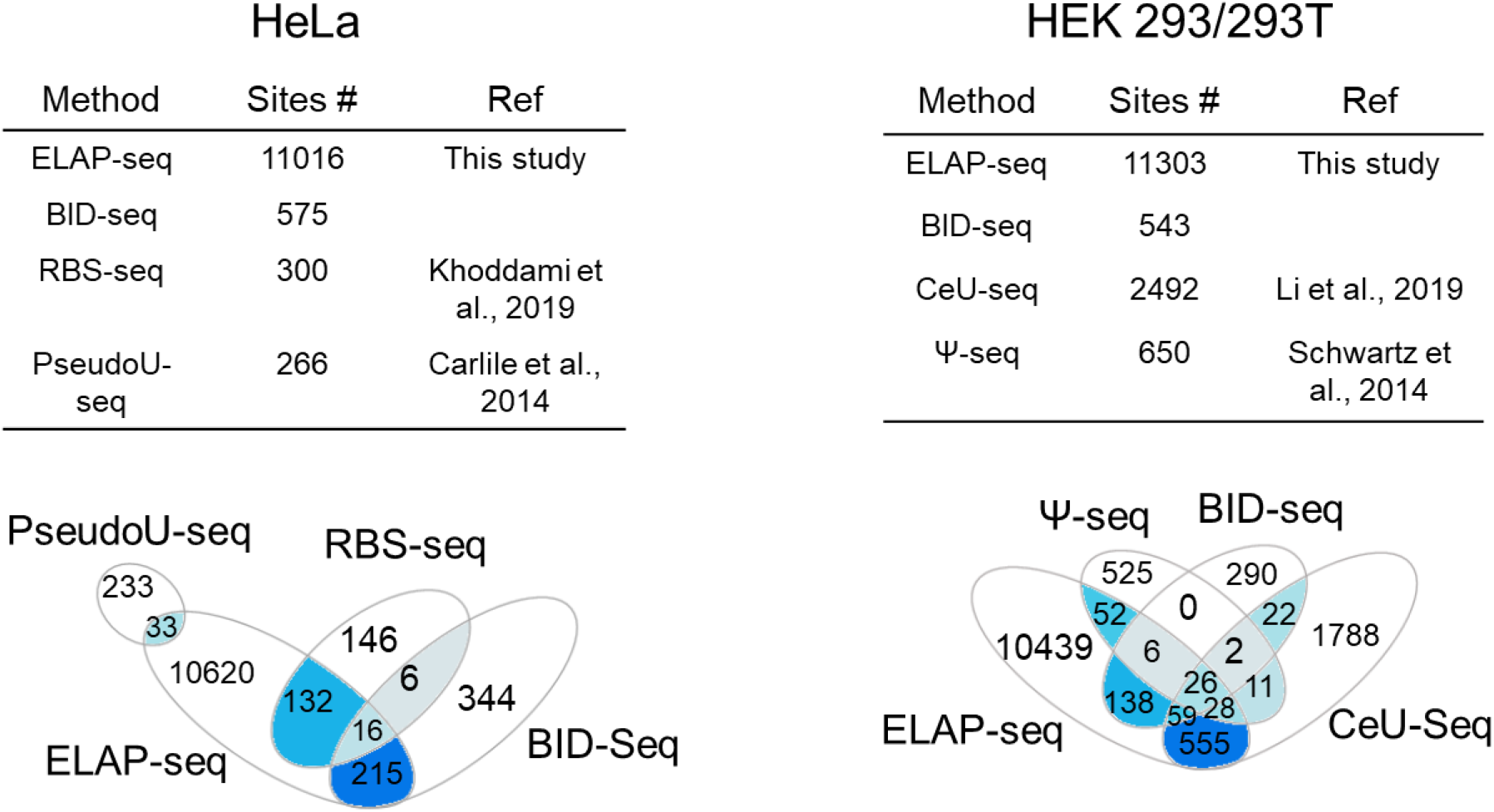
Overlap of sites identified by ELAP-seq and sited identified by previous methods.

**Figure S9.**
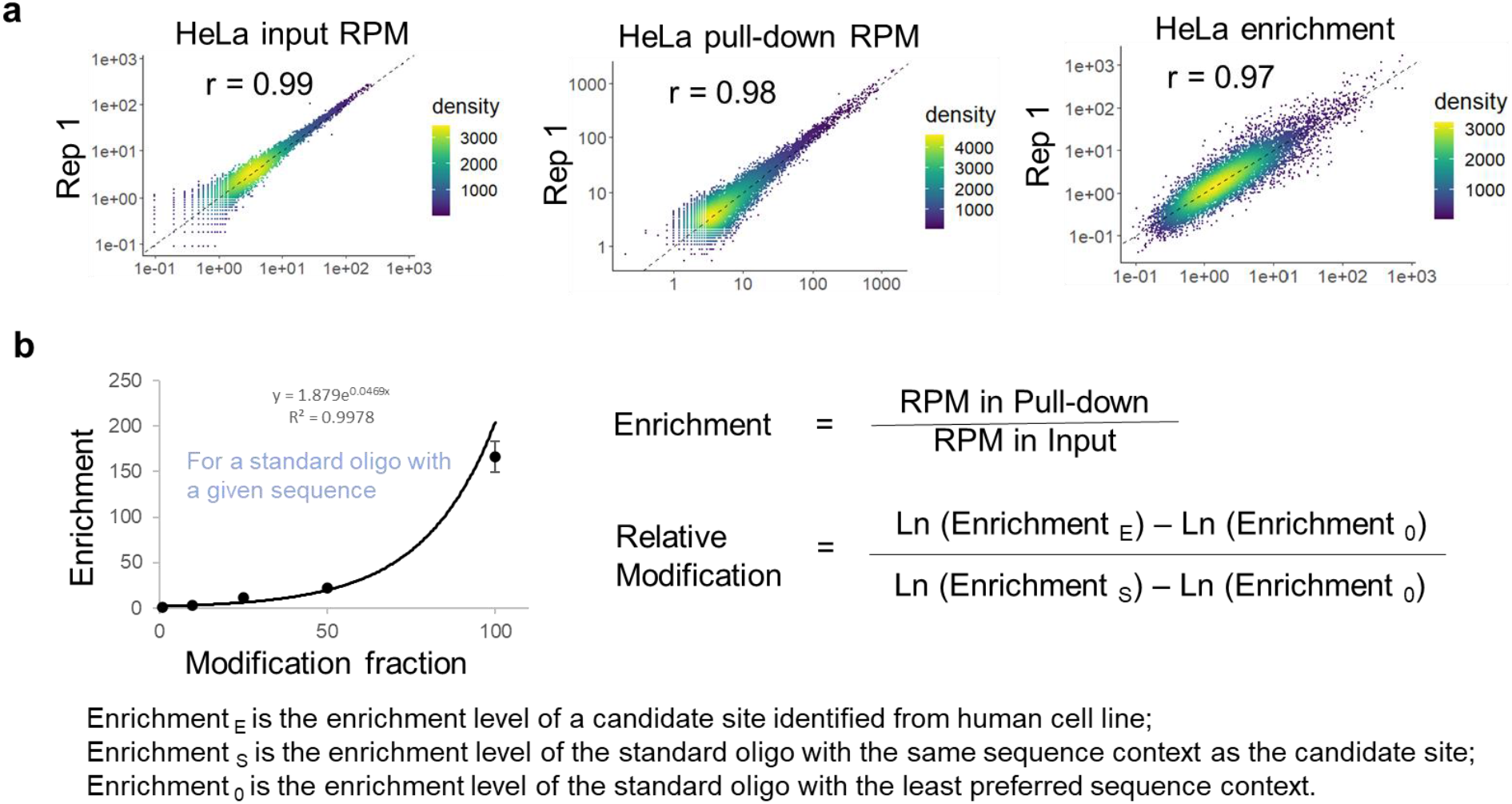
Parameters for semi-quantitatively evaluation of Ψ modification levels. a) RPMs in the input and pull-down samples, and enrichment levels are highly correlated between two biological replicates in both HeLa and HEK 293T cell lines. b) Equation deduced to semi-quantitatively evaluate relative modification levels of various sites based on the pull-down-qPCR result of a synthetic RNA and taking into consideration sequence context preference of the labeling strategy.

**Figure S10.**
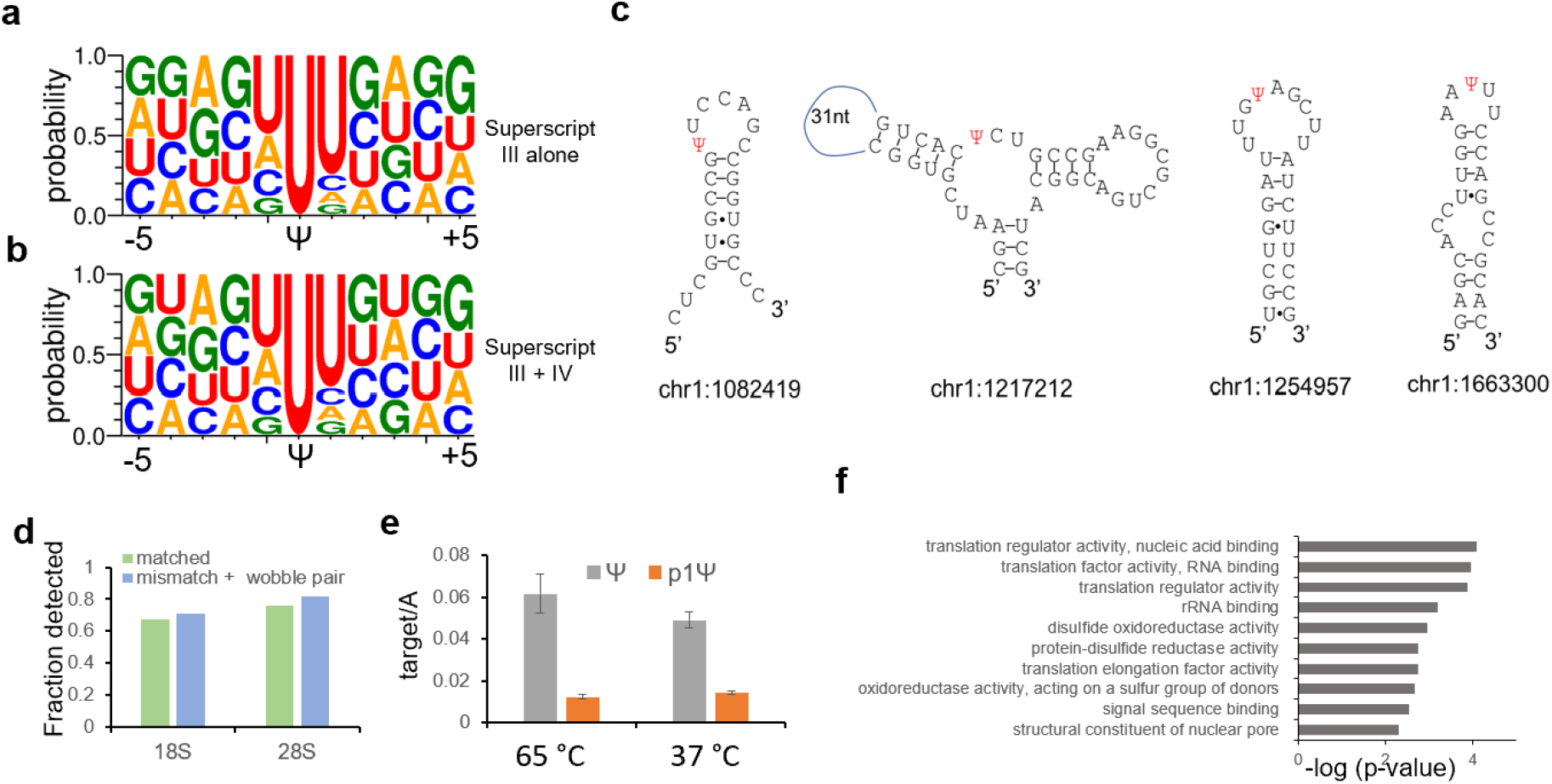
Implication on biological functions of Ψ modifications. a) Sequence motif of 11-nt surrounding the candidate Ψ sites of HeLa identified from libraries generated with Superscript III alone. b) Sequence motif of 11-nt surrounding the candidate Ψ sites of HeLa identified from combined sequencing reads of libraries generated with Superscript III and Superscript IV. c) RNA-fold analysis shows Ψ modification tends to locate in the less structured region. d) rRNA Ψ sites in matched regions and sites in mismatched plus wobble pair regions are equally detected by ELAP-seq. e) The enzymatic labeling efficiency on isolated total RNA at 37 °C is the same as that under the temperature of 65 °C. f) Top 10 enriched GO clusters of molecular functions for Ψ-modified genes in HeLa cells.

